# Intrinsic properties ensure reliable attractor dynamics in learned neural assemblies embedded within noisy, asynchronous networks

**DOI:** 10.1101/2022.07.26.501548

**Authors:** Matthieu Sarazin, Jeanne Barthélémy, David Medernach, Jérémie Naudé, Bruno Delord

**Affiliations:** Institut des Systèmes Intelligents et de Robotique, Sorbonne Université, Centre National de la Recherche Scientifique, UMR 7222, 75005 Paris, France; Institut de Génomique Fonctionnelle, Université de Montpellier, Centre National de la Recherche Scientifique, Institut national de la santé et de la recherche médicale, UMR 5203 CNRS, rue de la Cardonille 34094 Montpellier Cedex 05, France

**Author notes:** Equal contributions (Jérémie Naudé, Bruno Delord).

## Abstract

Neural representations rely on the ability of neuronal assemblies to display organized spiking patterns, despite being embedded within noisy networks. These structured patterns arise from attractor dynamics due to activity reverberation promoted by learnt assembly connectivity. Yet, attractor dynamics have been assessed either under low-noise conditions or in highly idealized neuronal assemblies. Here, in a spiking recurrent neural network model displaying asynchronous irregular noise, we show that realistic spike timing-dependent plasticity (STDP) imposes either low controllability of attractor recall (when STDP is strong), low stability of attractor maintenance (when STDP is low), or both. Moreover, STDP-built attractors display low independence, i.e., they perturb activity in the surrounding network. These constraints may favor self-generated representation switches essential for cognitive flexibility but dampening cognitive reliability. We reveal, by contrast, that several biophysical mechanisms alleviate these issues through a common dynamical principle, protecting excitatory-driven attractorial dynamics from inhibitory-driven spontaneous fluctuations. Amongst biophysical determinants, intrinsic properties were most efficient to increase controllability, stability and independence of attractor dynamics. Specifically, spike-triggered calcium-activated conductances improved reliability by mitigating reliance on connectivity, even at low conductance levels, i.e., in the absence of cell-autonomous intrinsic bistability. Finally, we show that the mechanisms we identify operate over a large repertoire of static (e.g., Hebbian or ring) and dynamic (e.g., sequences) attractors, with uni- and bidirectional propagation. Altogether, these results pinpoint synaptic and intrinsic synergy as a generic principle to regulate attractor reliability, as a function of the cognitive demand.

## Introduction

Neural trajectories — sequences of neural activity that propagate over time — represent local forms of ordered neural activity within the globally disordered awake brain regime. Neural trajectories act as dynamical codes across cerebral structures, supporting diverse cognitive operations, from motor function to working memory or navigation^1–6^. These trajectories do not simply mirror incoming time-structured sensory inputs: once triggered by a brief cue, neural trajectories can continue to propagate on their own.

Although many theoretical studies have examined activity sequences under simplified conditions for both background conditions and underlying synaptic connectivity ^7–17^, real neural activity is dominated by noise and connectivity is learned by synaptic plasticity.

Biological noise and learned connectivity introduce three critical constraints on neural trajectories that are often overlooked. First, random synaptic events can spontaneously trigger a neural trajectory, reducing its level of *controllability*, defined as the ability to trigger neural trajectories solely when needed. Second, noise may disrupt the *stability* of activity propagation once a neural trajectory has begun. While low levels of *controllability* and *stability* are desirable in the context of exploration and cognitive flexibility, much higher levels are expected for exploitation, cognitive control, reliable decision-making and deterministic volition. Third, a neural trajectory may interfere with other ongoing computations by strongly influencing nearby neurons, perturbing the *independence* of multiple simultaneous independent neural trajectories ^16^.

Understanding how biological circuits handle the levels of *controllability* and *stability* are to generate more or less reliable (i.e. controllable and stable) trajectories, depending on the cognitive demand, and how they overcome the problem of *independence*, is therefore a major objective for neuroscience.

Existing theoretical work has largely focused on mechanisms that stabilize static bumps of activity, for instance in ring-shaped networks, in strongly interconnected Hebbian assemblies of cells ^7,8^ or through cell-autonomous, intrinsic mechanisms ^18,19^. However, moving bumps (that model neural trajectories observed in the awake ^20–22^ or sleep ^23,24^ states) present additional difficulties. Models with idealized synaptic architectures, such as synfire chains (neurons connected along a feed-forward pathway within the network), asymmetric connectivity in ring models, or Hebbian phase sequences (HPS, i.e. oriented pathways between Hebbian assemblies) could produce moving bumps, but their stability under noise, their biological plausibility or their learnability is uncertain ^9,7,10,11,25,13^. Recurrent networks trained with artificial, biologically-implausible learning rules, can generate neural trajectories ^14,15,26^ yet these approaches do not clarify how real neural circuits might learn such connective pathways. A more biophysically-grounded mechanism for learning oriented connective pathways is spike-timing dependent plasticity (STDP) ^27^. STDP strengthens or weakens synapses based on the temporal difference between pre- and postsynaptic action potentials ^17,28^. However, the way trajectories created through STDP behave when embedded in a noisy recurrent network remains unresolved.

Most previous models based on idealized connectivity omit fundamental biophysical properties, such as distinct classes of synaptic receptors or voltage- and calcium-gated channels, whereas they represent major determinants of network dynamics in their ability to cast spiking patterns ^29,30^. The rich repertoire of intrinsic properties might offer powerful mechanisms for stabilizing trajectories, beyond the structure of the synaptic pathway itself. While intrinsic conductances have been proposed to help generate static bumps when strong conditions (i.e., thanks to neuronal bistability) are met^18,19^, it is unclear whether they can also support propagating sequences of neural activity.

Here, we present a dynamical and biophysical solution to control the level of reliability of neural trajectories in a detailed, data-driven cortical network operating in an asynchronous irregular regime. In our model, STDP and synaptic scaling produce a diversity of neural trajectories that differ in their stability, controllability and independence from surrounding chaotic dynamics. Neural trajectories emerge as a moving bump of activity driven by a strong, deterministic NMDA-mediated associative excitation at high firing frequencies. This contrasts to the network’s spontaneous state, dominated by low-frequency inhibitory GABA-A fluctuations. Using theoretical analysis, we identify a firing-frequency threshold separating spontaneous and propagating regimes. Mechanisms that increase the gap between these regimes improve both the controllability and stability of trajectories. We identify three classes of biophysical processes that widen such a gap: (1) low inhibitory fluctuations, (2) strong tonic feedforward inhibition combined with recurrent excitation, and spike-triggered calcium-dependent intrinsic currents that generate afterhyperpolarization and afterdepolarization. Among these, intrinsic currents provide the most robust and general solution, without requiring neuronal bistability. Spike-triggered calcium-dependent currents not only stabilize and control neural trajectories but also enhance their independence from the rest of the network, by reducing the reliance on synaptic mechanisms alone. These results are robust across model parameters and apply to many forms of neural trajectories — static, dynamic, discrete or continuous. In summary, we reveal a general dynamical principle — the coexistence of stochastic and deterministic regimes in the same network — and identify spike-triggered intrinsic currents as an efficient and plausible mechanism for learning and expressing neural trajectories with relevant levels of reliability, depending on the cognitive demand and despite the noise inherent to in vivo brain activity.

## Results

### Robust STDP-induced neural trajectories in the asynchronous irregular (AI) regime

Our goal was to understand how biophysical mechanisms help control locally ordered activity patterns — such as stable or moving bumps, REFs — that emerge from learned neural assemblies (REFs) embedded within globally noisy, vivo-like conditions (REFs) (**Fig. 1a**). Previous theoretical studies ^9–15,17,25,31–34,35^ showed attractor dynamics only in idealized network connectivity (REFs) or under low-noise conditions (REFs). Here we wanted a recurrent neural network model that was both noisy and with plausible connectivity, which would naturally emerge from synaptic plasticity applied to an initially random network.

**Figure 1.**
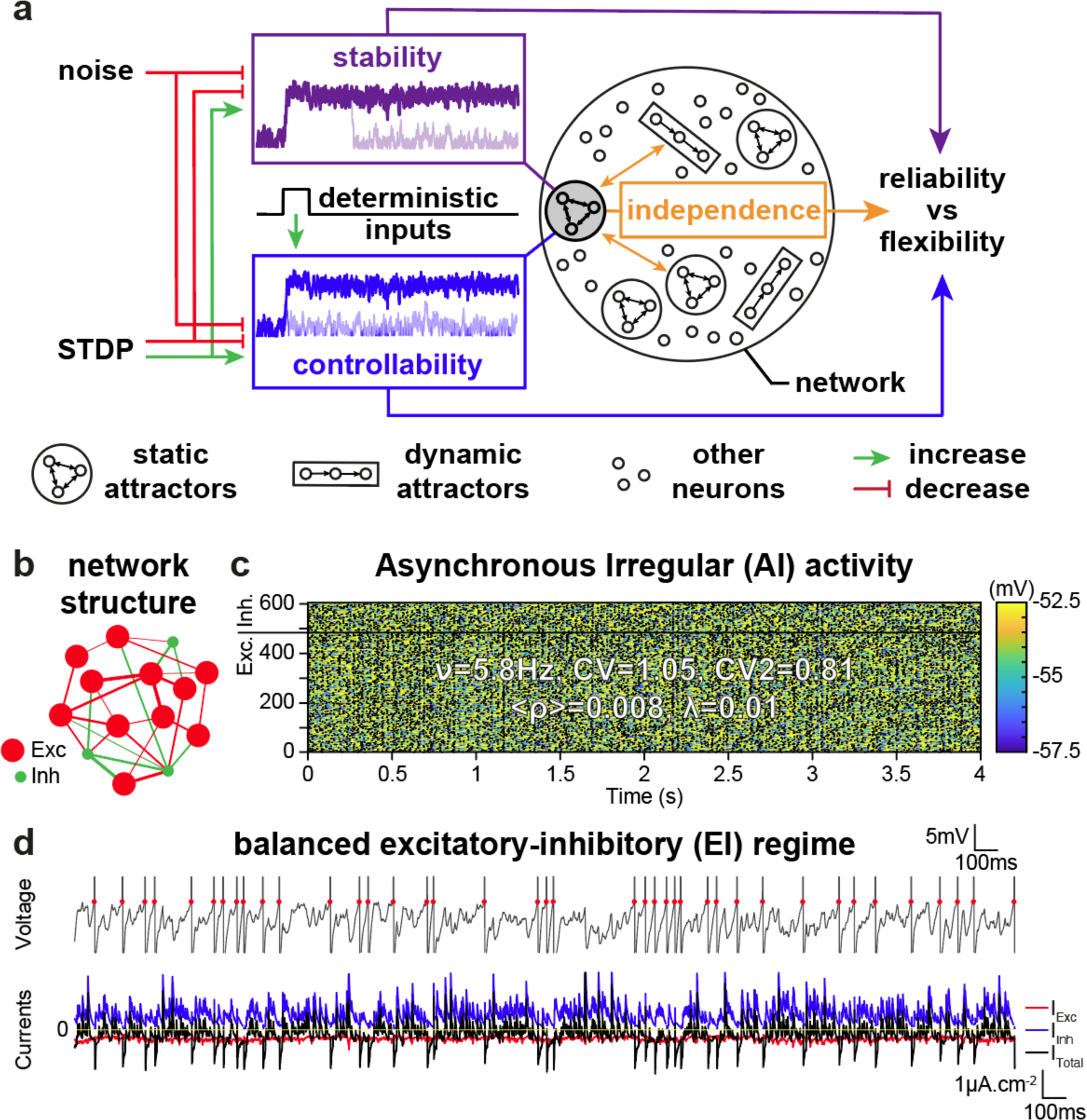
Properties related to reliability in a balanced spiking recurrent network. (**a**) Noise and spike timing-dependent plasticity (STDP) determine the degree to which activity in an attractor (shaded gray) 1) can be triggered – or not – by deterministic inputs (controllability, blue), 2) stably maintained – or not – once triggered (stability, purple), and 3) its degree of interference with other attractors or neurons in the surrounding network (independence, orange). Noise is globally deleterious to stability and controllability. STDP, by setting weights within attractors presents more complex effects that we study in the following. Altogether, increased levels of stability, controllability and independence favor the reliability of on-going computational processes, while their decrease promotes the flexibility of representations. (b) Randomly connected recurrent neural network of 80% excitatory and 20% inhibitory neurons (14 neurons are shown, whereas the model is composed of 605 neurons, i.e. 484 excitatory, 121 inhibitory). (**c**) Asynchronous irregular network activity, with spikes (black dots) and membrane potential of neurons across 4 seconds of simulation. (**c**) Subthreshold membrane potential and irregular spikes (top) driven by current fluctuations, since excitatory and inhibitory synaptic currents are balanced on average (bottom).

To keep the network model tractable ^36–38^ we avoided unnecessary complexities such as dendritic morphology or cortical layering (see *Methods*). We focused on biophysical mechanisms triggered by spikes: NMDA, AMPA and GABA synaptic currents, as well as CAN and AHP intrinsic currents (see *Methods*). Because these processes operate based on spikes, we used spiking integrate-and-fire models ^13,39^ rather than rate-based ^7,31^ or mean-field ^10,25^ models. We did not include detailed action-potential conductances, which slow down simulations and are not required for modeling the spiked-triggered processes of interest (see *Methods*). Finally, because the cortex exhibits both strong noise and organized neural trajectories^1–3,5^, we constrained the initial random connectivity using statistics from cortical data ^40–44^. Importantly, we did not impose any built-in structure such as ring attractors^7,10,11,25^ or pre-defined sequences of Hebbian assemblies ^13^.

We first verified that our model produced the hallmarks of cortical spiking activity (**Fig. 1b**, see *Methods*), (1) low firing frequencies (**Fig. 1c**, *v* < 10*Hz*, **Fig. S1b** ^45^), (2) high irregularity (inter-spike intervals (ISIs) *CV*∼1 and *CV*_2_∼0.8, and *LV*, described in *Methods)* **Fig. S1c-d** ^46^) and (3) asynchrony (average pairwise correlation < *ρ* > ∼0.01, **Fig. S2b-g** ^47^), all characteristic of the asynchronous-irregular regime ^39^ observed in behaving mammals ^48^. Accordingly, the network displayed chaotic dynamics (*λ*∼0.01, **Fig. S2e-j** ^49^), arising from fluctuation-based spiking (**Fig. 1d** top). This reflects a balanced “high-conductance” state^50^, in which strong excitatory and inhibitory currents track each other (**Fig. 1d** bottom, and **Fig. S1k** ^26^).

We next tested whether such a noisy network could learn a structured pathway through synaptic plasticity. Neurons were arranged on a 2D topographic map and we trained the network with a stimulus following a circular trajectory (**Fig. 2a**, left). Learning emerged from two plausible plasticity mechanisms, Spike-Timing Dependent Plasticity (STDP) and synaptic scaling (see *Methods*), which are ubiquitous in the neocortex ^51,52^. Synapses were potentiated when postsynaptic followed presynaptic spikes (**Fig. 2a**, center). As a result, sequential neuronal activation potentiated synapses along the direction of the stimulus, creating an oriented “*connective pathway”* i.e. an engram of the stimulus (arrows, **Fig. 2a**, right). Synaptic scaling stabilized learning by keeping the total incoming weights constant ^36,51^ (and thus similar across neurons, **Fig. 2a**, right).

**Figure 2.**
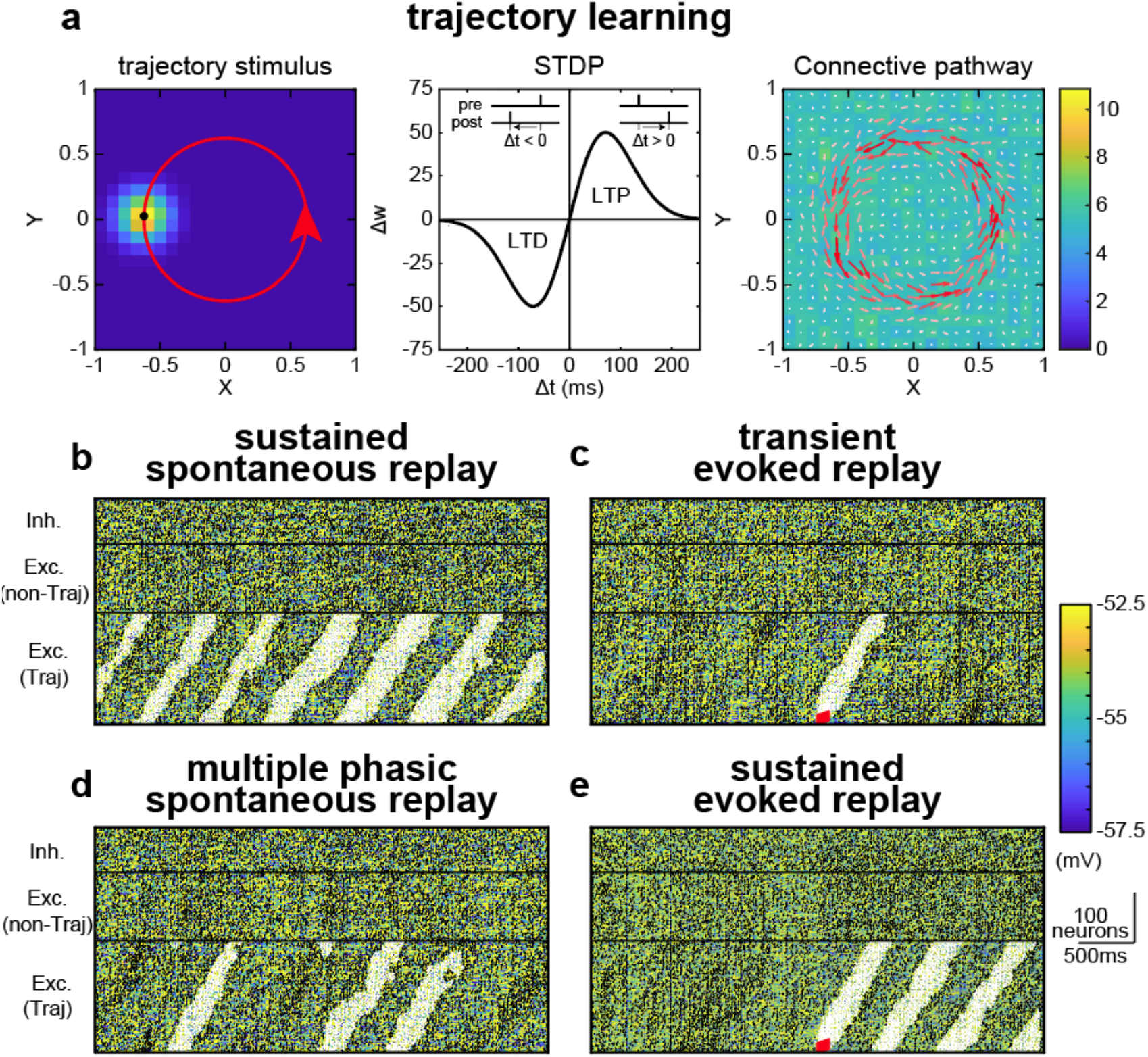
(**a**) (left) External circular trajectory stimulus (red circle), activating neurons through putative spatially-organized receptive fields. Example activity of neurons (background colors) induced by the trajectory stimulus at a given time point (black dot). (middle) Temporal window of the STDP learning rule, inducing LTP for positive time differences (pre-then post-synaptic spikes) and LTD for negative time differences (post-then pre-synaptic spikes). (right) Resulting connective pathway, with normalized arrows showing the direction in which outgoing weights are most potentiated (white to red arrow color scheme with increasing arrow magnitude), and homogeneous background colors showing similar sums of total incoming weights onto neurons (due to synaptic scaling) (**b-d**) Resulting connective pathways induce a variety of different trajectory replays, which emerge spontaneously (**b & d**) or can be evoked via a strong stimulus onto the first 25 neurons of the trajectory (red rectangle, **c & e**).

We next tested whether this learned connective pathway could propagate a neural trajectory (a *moving bump of activity)* defined as the set of co-active neurons within the engram (Methods). Neural trajectories in chaotic networks face distinct challenges: noise may disrupt bump propagation, so the connective pathway must be strong enough. But if synapses are too strong, random spikes can trigger uncontrolled bumps. Furthermore, bumps may spill over to surrounding neurons, disrupting other simultaneous neural trajectories. Indeed, after learning, excitatory activity amplified rapidly along the engram, recruiting nearby neurons and generating propagation involving ∼100 neurons, affecting global network dynamics (**Fig. 2b-e**).

Remarkably, the same network architecture could produce very different neural trajectories, depending on random initial connectivity before learning, or initial activity conditions — a consequence of chaotic dynamics. Spontaneous-persistent neural trajectories could emerge from noisy activity (**Fig. 2b, 2d**). Evoked but unstable trajectories could be triggered by an initiating stimulus but then fail to propagate (red rectangle, **Fig. 2c**). Evoked and stable trajectories could occur, displaying stable propagation when triggered (Fig. 2e). These patterns resemble a range of physiological observations such as free replay^20–23,53^ (i.e. low-rate spontaneous emergence and persistent **Fig. 2d**) or stimulus-triggered trajectories during correct task execution ^2,3,5^ (**Fig. 2e**). Thus, biologically realistic STDP can indeed train recurrent networks to express neural trajectories despite strong in vivo noise — but does not guarantee their reliability. To understand what makes trajectories reliable (stable, controllable, and independent) we next examined the dynamical conditions under which whether activity bumps do or do not propagate.

### Bump propagation relies on the transition from inhibitory fluctuation-to excitatory mean current-driven spiking

To understand how moving bumps propagate, we first examined what drives spiking in excitatory neurons depending on whether they belong to the moving bump (*“bump neurons*”) or not (“*non-bump neurons”*).

In *non-bump* neurons, the membrane potential settled on a subthreshold plateau (∼54mV, **Fig. 3a**, top left, arrow). The total current was near-balanced on average (**Fig. 3b**, top left, star). Spiking arose from fast voltage transients (**Fig. 3a**, top left, star) themselves caused by strong current fluctuations (**Fig. 3b**, bottom left, star). Among synaptic inputs, GABA-A inhibition fluctuated the most (**Fig. 3b**, bottom middle, star): brief drops in GABA-A current allowed the membrane potential to reach the threshold (**Fig. 3a**, bottom left, star). Thus, in *non-bump* neurons, spiking was mainly caused by disinhibitory fluctuations ^54,55^.

**Figure 3.**
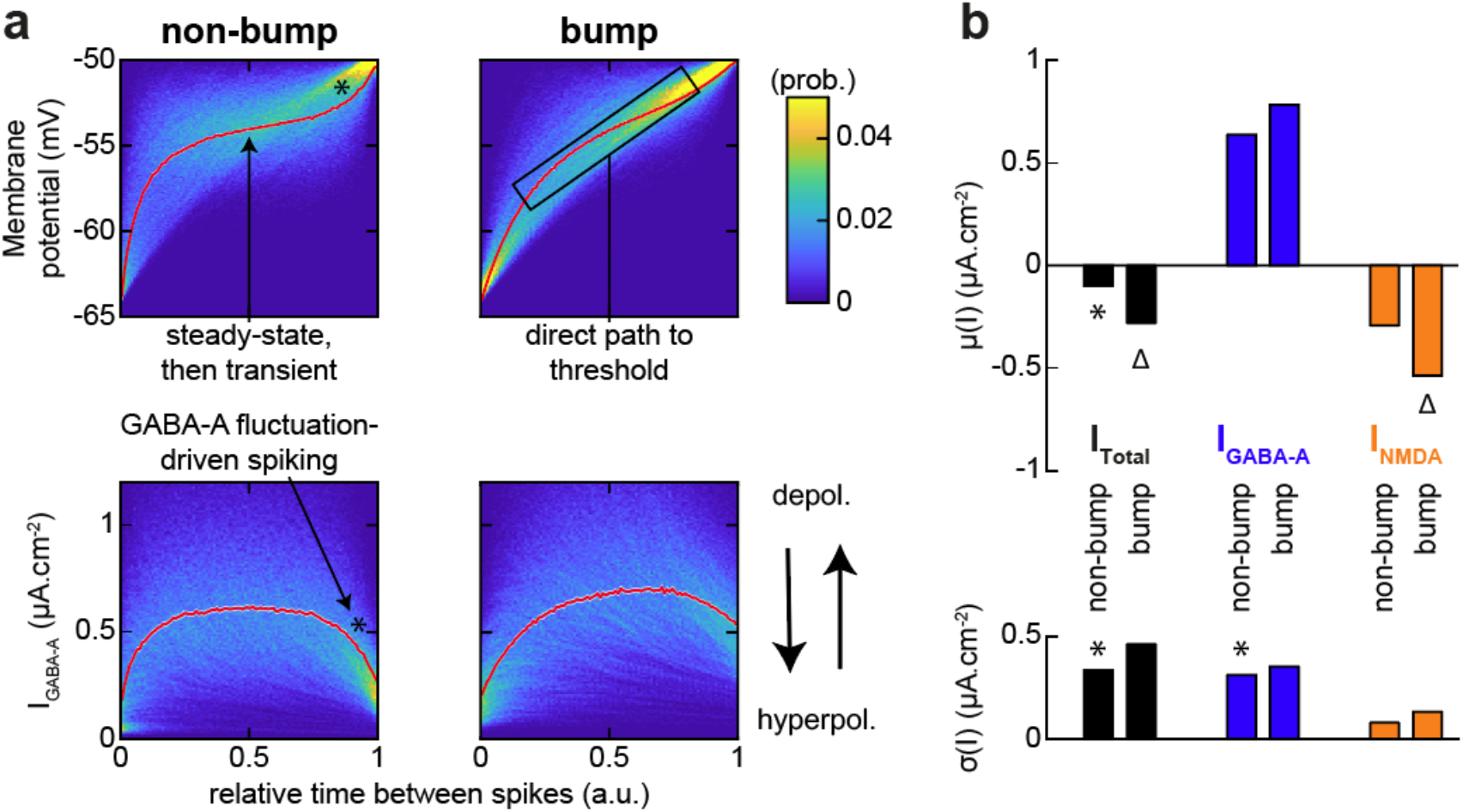
Bump propagation relies on a transition from GABA-A fluctuation-to NMDA mean current-driven deterministic spiking. (**a**) Membrane potential (top) and GABA-A current (bottom) of neurons when outside (left) and within (right) the trajectory activity bump, considered at the time scale of an ISI or between two spikes. Data is aggregated by normalizing time between two spikes (no matter the ISI duration). Background color shows the probability of individual membrane potentials or GABA-A currents curves (sum normalized to 1 in each time bin) across many ISI during one network simulation of 4 seconds, with red curves showing the average (weighted according to the underlying probabilities at each time bin). (**b**) Temporal average (top) and fluctuations (bottom) of total (left, black), GABA-A (middle, blue) and NMDA (right, orange) currents onto neurons when outside (non-Bump) or within (Bump) the trajectory activity bump, averaged across neurons.

In *bump* neurons, the membrane potential rose directly to threshold without passing through a plateau (**Fig. 3a**, top right). This happened because the total synaptic current was strongly depolarizing (**Fig. 3b**, top left, triangle), dominated by a strong, nearly-constant NMDA current (**Fig. 3b**, top right, triangle, and **Fig. 3a**, right). The NMDA drive was strong for three reasons: 1) bump neurons fired at a high frequency (*f*_*Bump*_∼14.5*Hz*), 2) synaptic weights along the connective pathway had been potentiated by STDP, and 3) coincident pre/post-synaptic activity removed the Mg^2+^ block of NMDA, resulting in “associative” NMDA activation among bump neurons. In addition, the bump triggered weak local inhibition, because inhibition was global (i.e. not specific to the bump), which further increased the imbalance between excitation and inhibition.

Overall, spontaneous global activity was dominated by GABA-A-fluctuations in a low-frequency, asynchronous-irregular regime^54,55^. By contrast, bump propagation relied on deterministic, associative NMDA excitation at high firing frequencies along the learned connective pathway. We next examined what controls the transitions between these two regimes, in order to identify the conditions under which neural trajectories start, propagate, or fail.

### Theoretical analysis of transitions between spontaneous activity and bump propagation

To study how activity switches between spontaneous activity and bump propagation, we first needed to identify where the transition between these two regimes occurs. Although global spiking noise defines the asynchronous-irregular state, the clearest marker distinguishing bump from non-bump neurons was their firing frequency (*f*), because firing rate determines the average synaptic currents and their fluctuations (**Fig. 3**).

We therefore measured the firing-frequency threshold separating regimes (**Fig. 4a**). To do this, we considered a simplified one-dimensional analytical model that predicts whether activity from a group of neurons propagate to the next ones along the trajectory pathway, as a function of *f* (numerically determined across neurons and network realizations for the standard “Model ∅”, see *Methods*). This simplified “*propagation-threshold model”* qualitatively predicted that this frequency threshold *f*_*θ*_ (**Fig. 4a**, left) is an unstable fixed-point: above the threshold, activity self-amplifies and propagates, while below the threshold, activity extinguishes. The *propagation-threshold model* reproduced average currents in spontaneous vs propagation regimes in the network model (**Fig. 4a**, right); as well as the threshold value itself (*f*_*θ*_∼10.2*Hz* – vs *f*_*θ*_∼9.2*Hz* in network simulations, see *Methods*).

**Figure 4.**
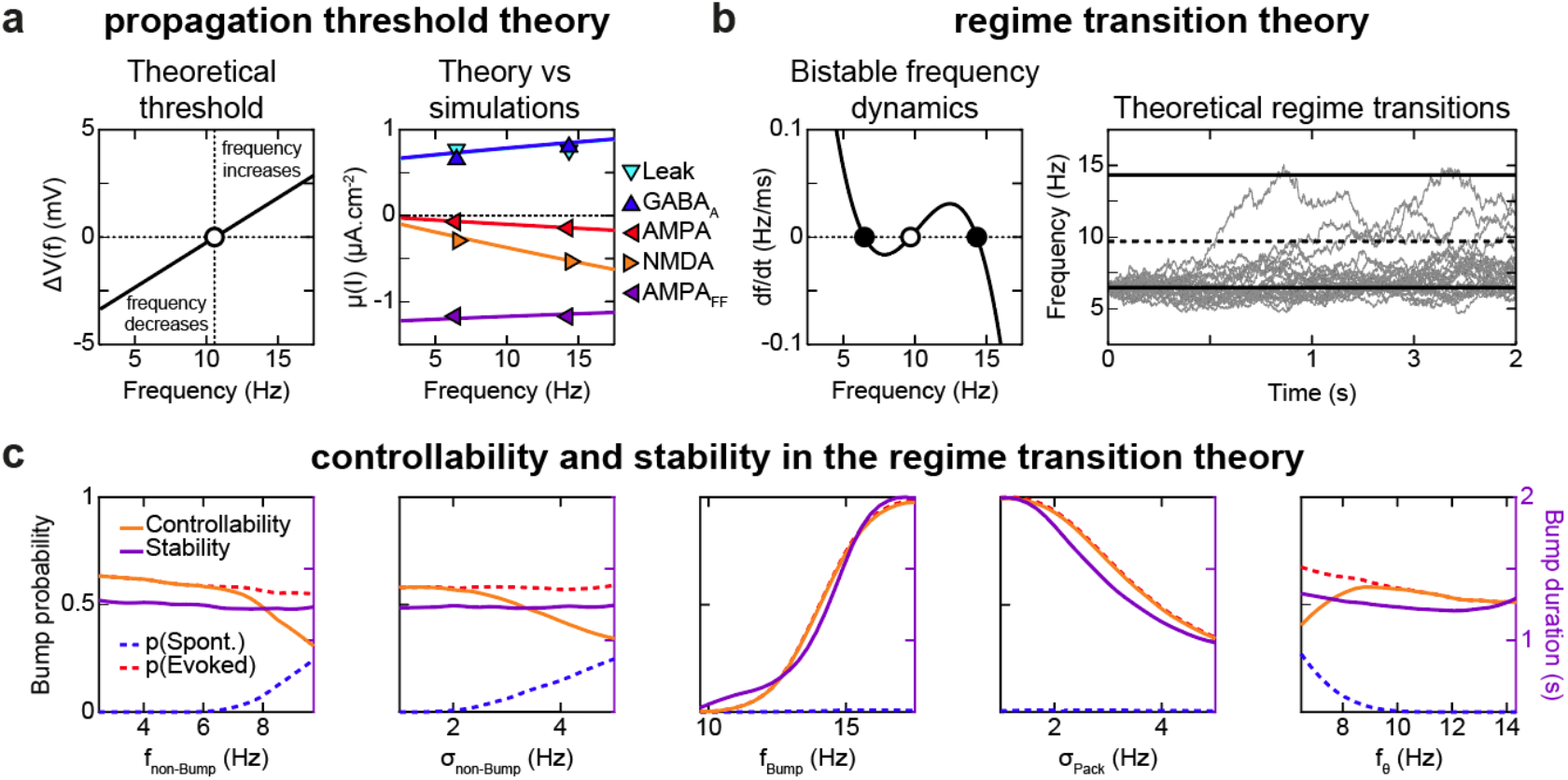
Theoretical account of the threshold separating, and transitions between, spontaneous and propagation regimes. (**a**) Propagation threshold model. (left) In a 2a reduced analytical model (see Methods), frequency self-amplifies above *f*_*>θ*_ and is extinguished below, i.e. as the membrane potential reaches (*ΔV* > 0) or not (*ΔV* < 0) a fluctuation-based spiking threshold at time *T* = 1/f in a postsynaptic neuron, given presynaptic spiking at frequency *f*. (right) The theoretical model (lines) is quantitatively consistent with network simulations ( symbols) at the fine-grain of ionic and synaptic currents in both the low frequency spontaneous and the higher frequency bump regimes. (**b**) Regime transition model. (left) In a 2a reduced model of both regimes, frequency dynamics follows bistable dynamics with added Gaussian noise. (right) Example simulations of the regime transition model. (**c**) Probability of spontaneous and evoked transitions to the regime of bump propagation (dotted lines), and of controllability and stability of the bump regime (solid lines) in the regime transition model, as a function of parameters (non-Bump and Bump mean frequencies and standard deviations, and the threshold frequency).

Based on the *propagation-threshold model*, we designed another simplified *regime transition model* (**Fig. 4b**, see *Methods*) to qualitatively predict how biophysical factors affect noise-driven transitions between the two regimes. We considered that the recurrent network can be summarized four key dynamical features: 1) a stable, low-frequency state (spontaneous activity), 2) a stable, high-frequency state (bump propagation), 3) an unstable threshold in between, and 4) stochastic firing-frequency fluctuations due to chaotic asynchronous-irregular dynamics. The simplest mathematical description capturing these features is a one-dimensional cubic differential equation. This *regime transition model* naturally produces two stable fixed points (*f*_*non−Bump*_ and *f*_*Bump*_) and one unstable point (the threshold *f*_*θ*_, **Fig. 4b**, left), plus a noise term that can induce transitions between states (**Fig. 4b**, right). The noise term, i.e. standard deviations *σ*_*non-Bump*_ and *σ*_*Bump*_, were estimated from network simulations.

The *regime transition model* allowed us to compute the probabilities of spontaneous *p*(*Spont*. ) and triggered *p*(*Evoked*) transitions toward bump propagation. We defined the *controllability* = (*p*(*Evoked*) − *p*(*Spont*. )) which captures the ability to trigger trajectories by the stimulus compared to spontaneous ones. To compare with numerical estimates in the full recurrent network, we simplify defined *bump stability* as the duration of evoked bump propagation (see *Methods*) rather than high–dimensional attractor (structural or marginal) stability measures ^15,56,57^.

From the simplified model, several qualitative principles for increased controllability emerged (**Fig. 4c**): 1) low firing rate or low firing variability in the spontaneous regime (*f*_*non-Bump*_ or *σ*_*non-Bump*_) decreased *p*(*Spont*. ) without affecting *p*(*Evoked*); 2) larger firing frequency (*f*_*Bump*_) or lower frequency variability (*σ*_*Bump*_) in the propagation regime increased *p*(*Evoked*), without affecting *p*(*Spont*. ); 3) controllability is maximal at intermediate *f*_*θ*_ that initially decreases p(*Spont*. ) but eventually p(*Evoked*) as well.

In summary, the simplified model predicts that biophysical mechanisms that increase the separation between the two regimes -either by pushing the bump firing rate higher, by lowering the spontaneous firing rate, or by reducing variability-should increase controllability. Furthermore, stability should increase when downward transitions from the propagation regime are decreased (i.e. higher *f*_*Bump*_ or lower *σ*_*Bump*_, **Fig. 4c**, third and fourth panels).

### Biophysical mechanisms promoting stability and controllability

The qualitative predictions from the simplified model guided us toward identifying the biophysical mechanisms (network architecture, synaptic and intrinsic currents) that affect reliable propagation in the full network model. To do this, we conducted extensive parametric explorations to find biophysical mechanisms that create a clear separation between the two activity regimes, ensuring both stability (bump propagation) and controllability (low probability of spontaneous bumps).

A straightforward approach was to strengthen the connectivity within the pathway by increasing STDP amplitude (*A*_*STDP*_). This enhanced stability because it raised the bump firing frequency (**Fig. 5a**, left). However, it also increased the probability of spontaneous bumps because *f*_*θ*_ shifted downward (**Fig. 5a**, left). As a result, controllability improved only marginally (**Fig. 5a**, middle).

**Figure 5.**
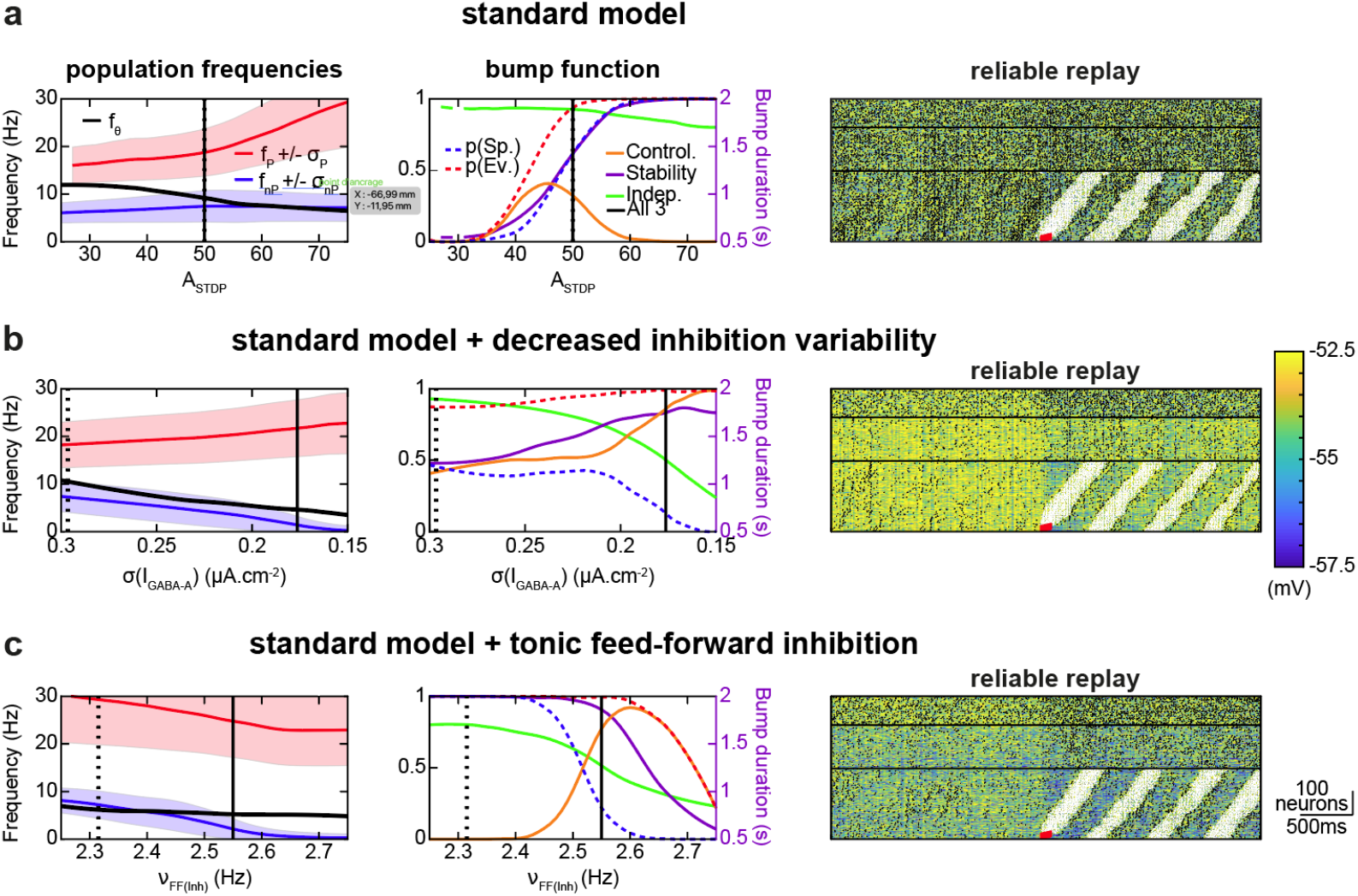
Modulation of trajectory control by architectural and synaptic mechanisms. (**a-c**) Mechanisms for increasing trajectory replay controllability and stability, compared to the standard model (**a**),under increased GABA-A current fluctuations *σ*(*I*_*GABA-A*_) (via a higher number of In->Exc synapses, see *Methods*) (**b**), higher AMPA feedforward currents onto inhibitory neurons *FF*_*Inh*_ (**c**). STDP amplitude *A*_*STDP*_ was varied across the different mechanisms (*A*_*STDP*_ = 47.5 for *σ*(*I*_*GABA-A*_), 75 for *FF*_*Inh*_), . (left) Non-Bump and Bump average frequency (+/-fluctuations) and threshold frequency, when varying the aforementioned parameters (X-axis). Normal parameter values (dotted vertical black lines), and those chosen to illustrate the mechanism’s effects on trajectory replay (solid vertical black lines), are indicated. (middle) Probability of spontaneous and evoked bumps (dotted lines), and bump controllability, propagation stability and independence (solid lines). (right) Example of trajectory replay with the selected illustrative mechanism parameters. The (*A*_*STDP*_, mechanism parameter) value couple of each mechanism was systematically determined as that maximizing the product of controllability, stability and independence (all three being normalized between 0 and 1).

We then reasoned that reducing the fluctuations of inhibitory currents onto excitatory neurons -while keeping their mean constant-should selectively suppress spontaneous transitions. Indeed, the spontaneous regime is mostly driven by disinhibition, whereas the evoked regime is not. To reduce inhibitory variability, we increased the number of *I* → *E* synapses while reducing their individual strength (higher *p*_*I*→*E*_, see *Methods*). This effectively lowered GABA-A current fluctuations (*σ*(*I*_*GABA-A*_), **Fig. S3a**, top). As predicted, lower *σ*(*I*_*GABA-A*_) decreased the firing rate in neurons outside the bump *f*_*non-Bump*_ (and thus *σ*_*non-Bump*_, **Fig. 5b**, left), which in turn decreased *p*(*Spont*. ) (**Fig. 5b**, middle), thereby increasing controllability. Stability was also improved: lowering *f*_*non-Bump*_ reduced the excitatory drive onto inhibitory neurons, which then decreased inhibition onto bump neurons, increasing *f*_*Bump*_. A similar effect was observed when we used slower and proportionally weaker GABA-A currents (**Fig. S3a**, bottom and **Fig. S3b**).

We next explored a combination of STDP and inhibition mechanisms. As shown above, increasing excitatory weights within the learned pathway (*A*_*STDP*_ = 75) increased *f*_*Bump*_ and decreased *f*_*θ*_ without modifying *f*_*non-Bump*_ (**Fig. 5c**, left, black vertical line vs. **Fig. 5a**, left, black vertical line). This widened the separation between the two activity regimes but also brought *f*_*non-Bump*_ closer to *f*_*θ*_, leading to a high probability of spontaneous bumps *p*(*Spont*. ) (**Fig. 5c**, left and middle). It is possible to counteract this side-effect by increasing tonic feedforward (i.e. *v*_*FF*(*Inh*)_), which decreased both *f*_*non-Bump*_ and *f*_*Bump*_ by the same amount (while keeping *f*_*θ*_ unchanged, **Fig. 5c**, left). As a result, *p*(*Spont*. ) and *p*(*Evoked*) both decreased, but their difference (i.e. controllability) increased for intermediate values, and stability also improved. In summary, this mechanism improves controllability by adding frequency-independent inhibition that suppresses spontaneous firing frequency, while allowing frequency-dependent to dominate during propagation. The same principle could be implemented in many other biophysical ways, making it a general strategy for improving controllability of moving bumps. For example, decreasing the leak current of inhibitory neurons (*g*_*L*(*Inh*)_), or reducing total currents onto inhibitory neurons (*g*_*X*→*I*_), also increased inhibitory firing in the spontaneous regime, and thus produced similar effects (**Fig. S2c-d**).

Beyond synaptic and architectural factors, we found that two intrinsic currents were key determinants of controllability and stability: the calcium-activated non-specific cationic (CAN) and the after-hyperpolarization potassium (AHP). Both currents are activated by calcium entry (through voltage-dependent calcium channels), and thus both depend on firing frequency. We considered a combination of a slow AHP that saturates at low firing frequency, with a fast CAN that saturates only at high firing frequency. AHP thus dominates at low firing frequencies and hyperpolarization is favored, but at high frequencies, CAN dominates and produces more depolarization (**Fig. 6b**, left). Together, CAN and AHP produce an unstable fixed point at *f*′_*θ*_∼12.7Hz (**Fig. 6b**, left) at the single-cell level. Below this firing frequency, AHP suppresses activity and therefore prevents spontaneous bump initiation. Above *f*′_*θ*_, CAN amplifies activity, stabilizing bump propagation. This intrinsic mechanism greatly widened the separation between two regimes regimes at the network level, by lowering *f*_*non-Bump*_ and raising *f*_*Bump*_ (**Fig. 6a**, left). As a result, both controllability and stability were improved (**Fig. 6a**, middle). This effect did not require strong conductances that would generate full intrinsic bistability (i.e. persistent activity ^18,19^). A modest CAN conductance, sufficient to yield transient bistability, was enough to achieve strong controllability and stability of neural trajectories (**Fig. 6b**, middle).

**Figure 6.**
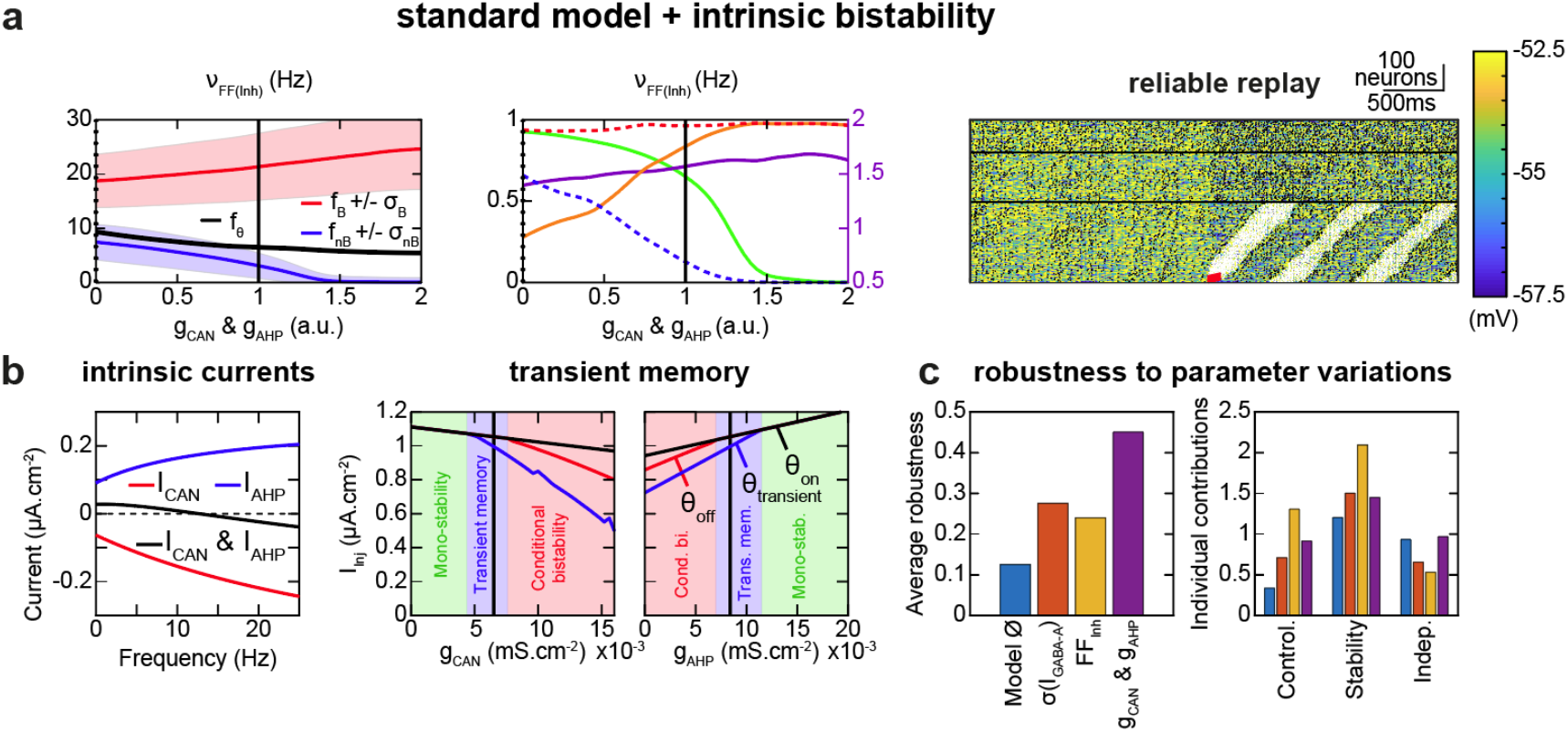
Improved trajectory control by intrinsic supra-threshold conductances. (**a**) Mechanisms for increasing trajectory control through increased CAN and AHP calcium-activated suprathreshold conductances together *g*_*CAN*_ &g_*AHP*_. Panels organized as in Fig. 5. (**b**) (left) Equilibrium values of CAN (red), AHP (blue) and total (CAN & AHP, black) currents of excitatory neurons, when considering the time-averaged calcium concentration at different spiking frequencies; (right) CAN and AHP calcium-activated suprathreshold conductances induced transient spiking bistability (rather than mono-stability, conditional bistability or absolute bistability), as defined by the protocol in ^38^ (see *Methods*). Solid vertical black lines indicate the chosen biophysical parameters. (**c**) (left) Average robustness of the physiological low-frequency asynchronous irregular network activity with controllable, stable and independent trajectory replays, to the variation of 22 of the model’s parameters (see *Methods* and **Fig. S3**). The standard model and its variations in Fig. 5 and 6a, and the standard Model (∅), are compared. (right) Contribution of the trajectory replay controllability, stability and independence criteria to the overall robustness score.

### Independence between trajectory and surrounding activity

For most biophysical mechanisms considered, improving controllability and stability required reducing spontaneous firing rate. But doing so creates a side effect: during bump propagation, the firing frequency of neurons outside the trajectory engram increases, leading to a sharp contrast between background network dynamics at rest and during neural trajectories (**Fig. 5b-c and Fig. 6a**, right). Such strong network-wide modulation might interfere with parallel computations, for example if multiple trajectories need to coexist simultaneously. To quantify these perturbations, we defined an *independence* measure 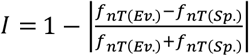, with *f*_*nT*_(*Ev*. ) and *f*_*nT*_(*Sp*. ) being the firing frequency of excitatory neurons outside the trajectory during bump propagation and during the spontaneous activity, respectively (**Fig. 5a-c and Fig. 6a**, middle). *I* = 1 means the two firing frequencies are identical, decreasing *I* means increasing difference, and *I* = 0 occurs when one of the frequencies is zero. In general, increasing controllability and stability decreased independence (down to ∼50%, from a ∼90% baseline). However, the CAN/AHP mechanism better preserved independence (down to ∼70%) by reducing the contrast between spontaneous and trajectory-related dynamics. These results show that combining synaptic factors (the learned connective pathway) with intrinsic properties (CAN/AHP)^38^ is computationally advantageous. CAN-AHP combination provides individual cells with a mild, transient tendency toward sustained firing (**Fig. 6b**, middle and right), thereby lessening the burden on synaptic pathways to maintain stable bump propagation.

### Robustness of reliable neural trajectories within the asynchronous-irregular regime

To test how general these mechanisms are, we evaluated how robust they remain under the wide biophysical variability observed across cerebral structures and species. We measured how large the parameter ranges are, in which both reliability (controllability, stability and independence) and asynchronous-irregular dynamics are preserved. We performed this robustness analysis by systematically varying parameters for each mechanism (see *Methods* and **Fig. S4a-c**).

In the standard model without any added reliability mechanism (Model ∅), controllable and table trajectories existed only within a moderately small parameter region (overall robustness ∼10%). With the additional mechanisms described above, these regions expanded substantially (∼25%), reaching a maximum with the CAN-AHP mechanism (∼45%, **Fig. 6c**, left).

However, the different reliability mechanisms did not affect independence similarly. Although the *v*_*FF*(*Inh*)_ mechanism resulted in the largest parametric regions for controllability and stability (when considered on its own), it reduced the parametric range for independence, as did *σ*(*I*_*GABA-A*_). Independence robustness could fall to ∼60% of the region width from the model ∅. In contrast,the CAN-AHP mechanism preserved ∼100% of the independence range, and further supported intrinsic currents as generic candidates for bump propagation alleviating constraints on synaptic-based propagation (**Fig. 6c**, right).

### A common framework for generalized static and dynamical neural attractors

Beyond robustness, genericity lies in the functional versatility. We thus evaluated whether the biophysical mechanisms for bump reliability could support the wide repertoire of attractor types described in the literature. Previous studies demonstrated many static and dynamic attractor types by using idealized engrams ^13^ or artificial training rules optimizing the connectivity ^15,31,32,12,58^. Here, we asked whether all these attractors could instead emerge with 1) activity-dependent plasticity rules and 2) the biophysical solutions for reliability identified above. To test this, we varied stimulus properties and STDP parameters (**Fig. S4a**), known to differ across cerebral structures ^52^ or neuromodulatory levels (e.g. dopamine ^59^). We then assessed to what extent the biophysical factors we uncovered (Fig. 4) support the controllability and stability of a broad variety of attractor types.

A static-discrete attractor is simply a stable non-moving bump of activity: a group of neurons that activate each other because they are strongly interconnected (visible as square blocks on the diagonal of the synaptic, **Fig. S4b**). This is the classical “Hebbian assembly” (HA). . We tested whether a symmetric STDP window (**Fig. 7a**, center and **Fig. 7b**, center), combined with discrete stimuli, could create such static-discrete attractors. A single discrete stimulus (**Fig. 7a**, left) formed of a single HA (**Fig. 7a**, right). When we added the CAN/AHP intrinsic mechanism, the stimulus reliably triggered persistent activity in the HA (**Fig. 7c**), showing that synaptic ^8,60,61^ and intrinsic ^18,19^ mechanisms can advantageously be combined to control static attractors. A stimulus that jumps between multiple discrete positions (**Fig. 7b**, left) produced multiple HAs (**Fig. 7b**, right). Thanks to the CAN/AHP mechanism, each HA could be independently triggered^62,63^ with higher reliability **(Fig. 7d**).

**Figure 7.**
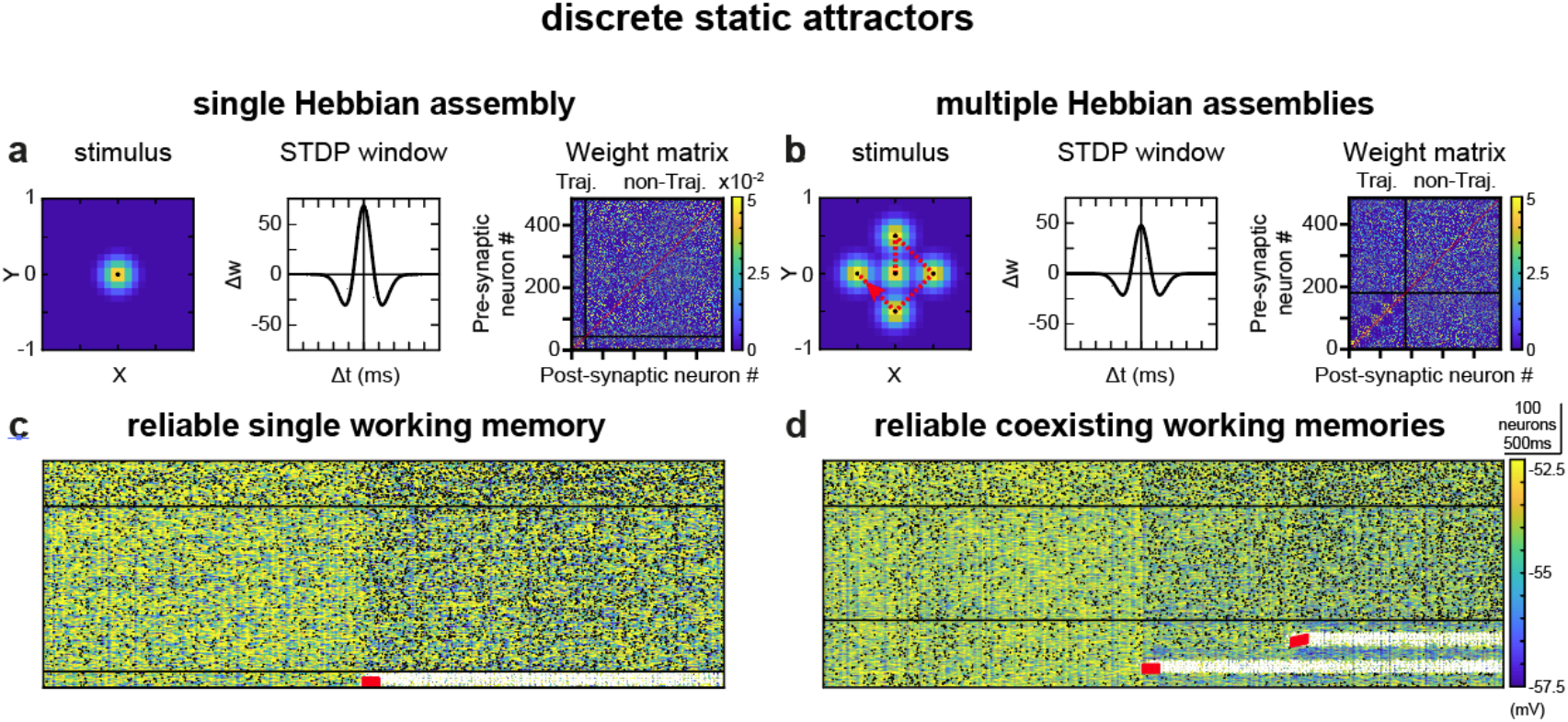
Discrete static attractor control with the intrinsic mechanism. In the case of discrete static attractors, i.e., single (**a**) or multiple (**b**) hebbian assemblies, increased supra-threshold conductances *g*_*CAN*_ & g_*AHP*_ express reliable single (**c**) or coexisting (**d**) persistent activity(ies), as found in working memory. In (**a**) and (**b**): (left) External trajectory stimulus (as in **Fig. 2.a** left). Dotted red lines indicate a discontinuous trajectory, jumping from one black dot to the next in a discrete manner (rather than continuously, as in **Fig. 9**). (middle) STDP temporal window (as in **Fig. 2.a** middle). (right) Resulting synaptic weight matrices between presynaptic and postsynaptic excitatory neurons. Neurons affected by the trajectory are regrouped and ordered according to their activation time within the learned trajectory stimulus.

Unlike static attractors, dynamical-discrete attractors involve activity that moves from one neuronal group to the next. A classic example is the synfire chain: activity propagates across neurons arranged in a feedforward sequence (off-diagonal synaptic patches, **Fig. S4c**). A series of HA can also be connected into a Hebbian Phase Sequence (HPS), where each assembly is strongly connected internally and to the next one (diagonal and off-diagonal patches, **Fig. S4d**). In HPS, the network activity can propagate from one HA to the next. We tested how an asymmetric STDP window combined with a sequence of discrete stimuli could generate these different dynamical-discrete attractors. A stimulus moving step-by-step (**Fig. 8a**, left), with a strongly asymmetric STDP window (**Fig. 8a**, center), produced a synfire chain^64^ (**Fig. 8c**) with fast, reliable propagation, supported by *σ*(*I*_*GABA-A*_) and *v*_*FF*(*Inh*)_ mechanisms, **Fig. 8c**). The same stimulus (**Fig. 8b**, left), learned with a weakly asymmetric STDP window (**Fig. 8b**, center) led to HAs linked by feedforward connections (**Fig. 8b**, right), i.e. a Hebbian phase sequence (ref 10) with slow sequential propagation between HAs (**Fig. 8d**). Hence, reliable synfire chains and HPS can emerge from STDP, without requiring hand-crafted connectivity matrices or artificial training rules ^13,64^.

**Figure 8.**
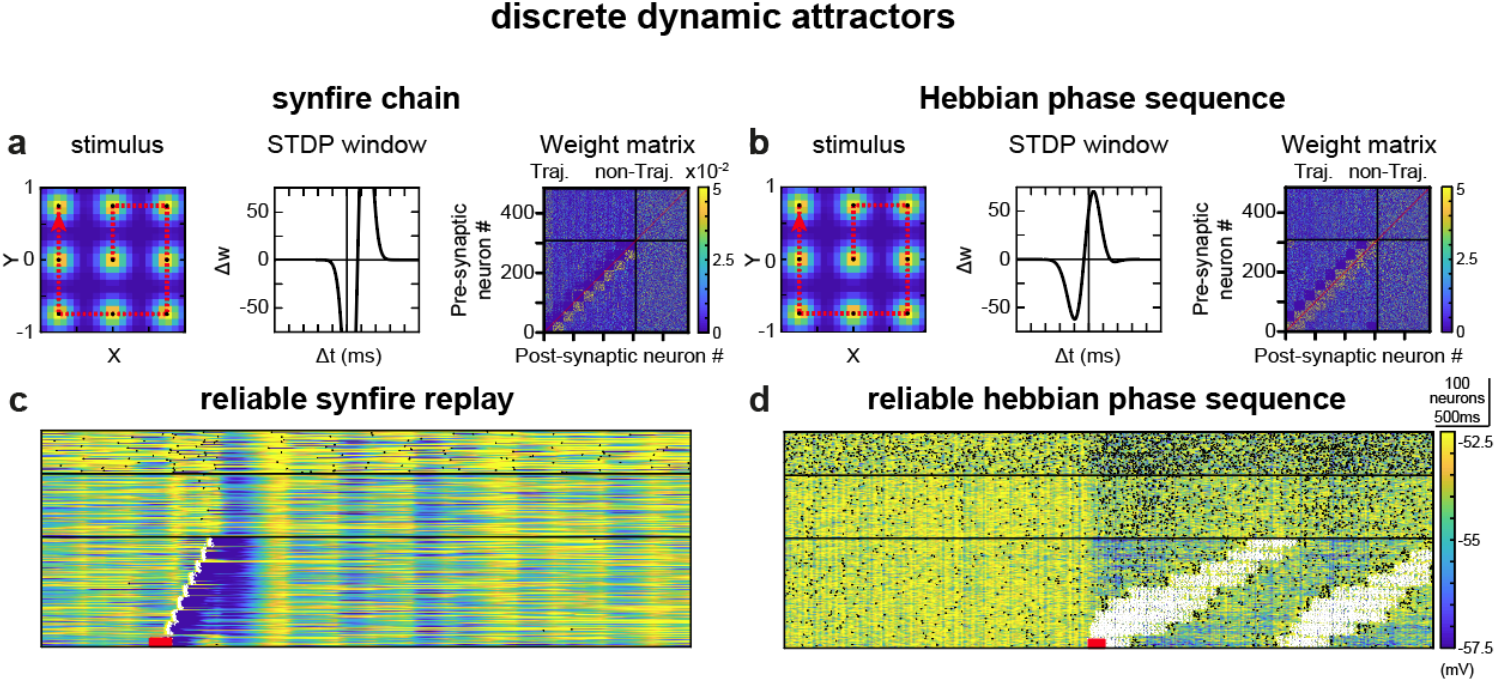
Discrete dynamic attractor control with synaptic mechanisms. In the case of discrete dynamic attractors, i.e., synfire chains (**a**) or Hebbian phase sequences (**b**) architectures, the *σ*(*I*_*GABA-A*_) mechanisms express reliable propagations of activity (**c, d**). Additional modifications were necessary for the synfire chain (**a, c**): the use of the *v*_*FF*(*Inh*)_ mechanism, *g*_*AMPA*_ = 0.5 *mS. cm*^−2^, g_*NMDA*_ = 0 *mS. cm*^−2^ instead of 0.2 *mS. cm*^−2^, g_*NMDA*_ = 0.3 *mS. cm*^−2^ for rapid bump propagation. (**a**) and (**c**): same organisation as in Fig. **7a, b**. Dotted red lines indicate a discontinuous trajectory (as in **Fig. 8**), rather than a continuous one (as in **Fig. 9**).

Unlike discrete attractors, continuous attractors allow activity to occupy a continuum of positions, and may also be static (fixed bump) or dynamic (moving bump). A classical example of continuous-static attractor is the ring attractor (REFS), where neurons are symmetrically connected to nearby neighbors (a diagonal band of strong weights, **Fig. S4e**). This supports static bumps anywhere on the ring. In our terminology, neural trajectories (the main focus above) corresponds to continuous-dynamic attractors. Neural trajectories can propagate unidirectionally or bidirectionally, depending on whether neighboring neurons are connected asymmetrically or symmetrically (off-diagonal bands, **Fig. S4f**).

We obtained a ring-like attractor (**Fig. 9a**, right; ^10,11,65^) by presenting a continuously moving stimulus (**Fig. 9a**, left) with symmetric STDP (**Fig. 9a**, center). Multiple neural trajectories could coexist, and drifted very slowly thanks to the CAN-AHP mechanism (**Fig. 9c**). Using the same stimulus (**Fig. 9b**, left) but with a broader STDP window (**Fig. 9b**, center) produced a wider symmetric connectivity (**Fig. 9b**, right). Combined with slowly-saturating *g*_*AHP*_ (see *Methods*, **Fig. 9d**), the network expressed bidirectional neural trajectories.

**Figure 9.**
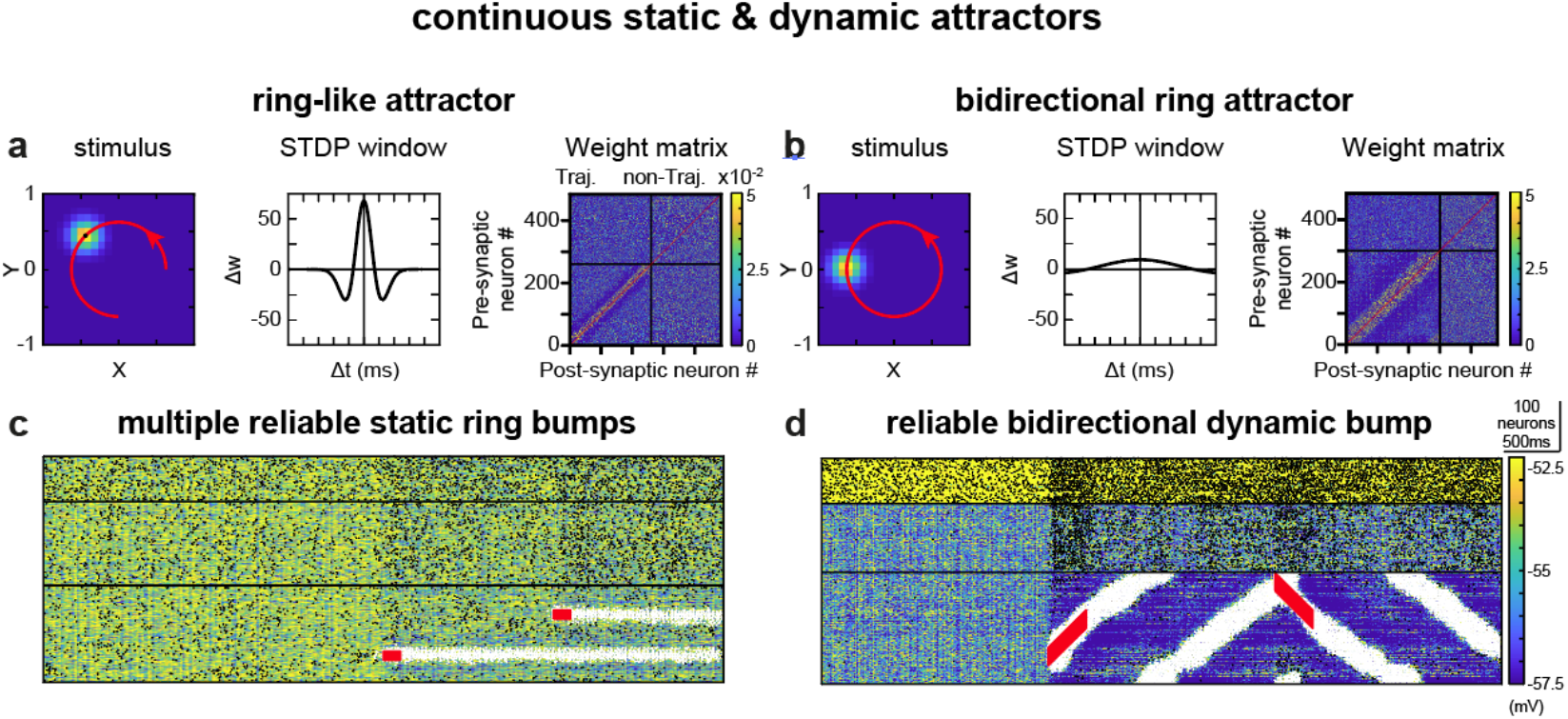
Continuous static and dynamic attractor control with intrinsic and synaptic mechanisms. In the case of continuous static and dynamic attractors, i.e., static (**a**) and bidirectional (**b**) ring architectures, the *g*_*CAN*_ & g_*AHP*_ and *σ*(*I*_*GABA-A*_) mechanisms, respectively, express reliable maintenance of static (**c**) or bidirectionally propagation (**d**) of bumps of activity. Additional modifications were necessary for the bidirectional ring architecture (**b, d**): *g*_*AMPA*_ = 0.5 *mS. cm*^−2^, g_NMDA_ = 0 *mS. cm*^−2^ instead of 0.2 *mS. cm*^−2^, g_*NMDA*_ = 0.3 *mS. cm*^−2^ for rapid bump propagation. (**a**) and (**b**): same organisation as in Fig. **7a, b**. Dotted red lines indicate a continuous trajectory (rather than discontinuous, as in **Fig. 7** and **Fig. 8**).

Altogether, these results show that the mechanisms we identified - both synaptic and intrinsic-provide general dynamical tools that allow the network to learn, stabilize and control a wide variety of dynamical representations. These include many classical attractors proposed for cognitive functions, suggesting a unifying biophysical framework for their emergence.

## Discussion

Using simulations and theoretical analyses of biophysically-constrained network models, we show how the interplay between intrinsic neuronal properties and synaptic connectivity enables recurrent neural networks to robustly control the reliability of attractor dynamics, including neural trajectories. These attractor dynamics persist despite the permanent disruptive influence of neural noise under the chaotic asynchronous irregular (AI) dynamics typical of wakefulness. Unlike classical approaches that focus on idealized synaptic connectivity, our work highlights the crucial role of intrinsic biophysical mechanisms - especially calcium-dependent conductances driven by spiking activity-in supporting the reliable emergence and maintenance of neural trajectories.

### Models of learning neural trajectories

Neural trajectories are widespread in the brain and are key to many cognitive operations ^5,21,22,66^. Theoretical models have proposed several mechanisms for how these neural trajectories might arise and remain stable ^13–15,17,28,33^. Here, we focused on three key physiological features that are usually studied separately but not together: 1) the destabilizing effect of noise due to asynchronous irregular (AI) dynamics and 2) synaptic connectivity learned by plausible Spike-Timing Dependent Plasticity (STDP).

Most STDP-based models do not test whether learned trajectories remain reliable under realistic neural noise^17,28^. Other models avoid these issues by using fixed, hard-crafted connectivity patterns (such as Hebbian phase sequence^13^), non-biological (non-local) training rules^14,15,31,58^, or by assuming unrealistically quiet neural activity^12,32,67^. In vivo, however, noisy fluctuations can interact with synaptic learning in complex ways that may disrupt memory dynamics or generate pathological activity ^36,68,69^.

We therefore examined how noise affects neural trajectories in the asynchronous-irregular state, where spiking activity emerges from a balance between excitation and inhibition^54,55^. The asynchronous-irregular state in our model relates to the classical “spike” chaos - with relatively constant spiking rate - and may differ from an heterogenous form of asynchronous-irregular state under strong synaptic coupling, with variable spiking rates across time and neurons (“rate” chaos ^70,71^). Consistent with previous work^36,69^, we found that spike chaos can undermine attractor stability, but we further identify specific biophysical mechanisms that allow recurrent networks to trigger and maintain neural trajectories in a plausible biological setting - connectivity learned by a STDP rule and under asynchronous-irregular noise conditions.

### Models of bump stability and propagation

Persistent “bumps” of neural activity have been studied extensively as low-dimensional attractors in recurrent networks. Classical bump models assume an idealized ring-like connectivity pattern with local excitation and surrounding inhibition^10,11,65^. This “Mexican-hat” architecture is anatomically unrealistic and is not used in our study^72^. Instead, bumps in our model arise from a synaptic engram embedded within a random recurrent connectivity that includes significant weights outside the engram (compared to usual negligible weights^13^). Disinhibition plays a different role here than in classical models: because inhibition is globally dominant in the AI regime (as in ^31^), fluctuations in inhibitory input mainly determine when non-bump neurons fire. Our work also integrates interactions between engrams, recurrent connectivity, feed-forward inputs^31^, synaptic scaling ^73^ and intrinsic properties^34^ - elements that have been treated separately in earlier models. We confirm that the strong inhibitory feedback used in formal models^39,74^ is also needed for static bumps (e.g., working memory) in the more realistic situation modeled here. For moving bumps, our results instead highlight the importance of strong external excitation onto inhibitory interneurons yet with small inhibitory fluctuations. Unlike earlier models that require asymmetric^10,25^ or complex^13^ connectivity to produce unidirectional propagation of bumps, our results support the possibility of bidirectional propagation under symmetric connectivity, which is absent in symmetric ring models ^7,25^ or in ^31^. We also show that bump stability can be maintained despite heterogeneous synaptic weights ^31,34,57,65,73^ (but see ^10^).

### Relation to previous neural reduced models

A major contribution of our work is identifying that the interaction between intrinsic and synaptic currents can support attractor reliability, i.e., controlability, stability and propagation of bumps. This is demonstrated through detailed simulations of the full model and in the low-dimensional simplified model. For the design of our simplified models, we followed classical strategies aimed at capturing essential dynamical features of the network. We used average firing as an effective measure to distinguish neurons currently inside the moving bump from those outside it. In the propagation model, average firing frequency of neurons is linked to how quickly membrane potential returns to baseline following spike repolarization, as classically done, in another context for theoretical frequency-intensity curves^75^. For the regime transition model, we used a one-dimensional cubic dynamical system, which is canonical to model two stable fixed-points separated by an unstable one. Such landscape is classical in computational neurosciences and explains molecular memory^76^, excitability-based bistability^77^ or working memory^78^. Yet, in our simplified models, we could relate spontaneous and propagation regimes (Fig. 2) and their frequency threshold (Fig. 3) with underlying biophysical mechanisms, contrasting with purely phenomenological rate models^79,78^. This allows us to link bump stability propagation and regime transitions to biophysical mechanisms ( synaptic and ionic currents, conductances, gating variables, and time constants) that shape network dynamics.

The reduced model was devised as a heuristic to identify which biophysical parameters are likely to influence bump propagation. It was not intended to quantitatively match all the details from the full recurrent dynamics – not to mention the large parameter exploration we performed. Instead, the simplified model offers qualitative predictions about how different biophysical mechanisms affect bump propagation and regime separation.

### Reliable trajectories beyond synaptic connectivity

Although the full repertoire of biophysical properties has been shaped by evolution to confer neural circuits with solutions to cognitive demands, computational models usually focus on synaptic connectivity alone^13–15,17,33,39^. Theoretical work on neural trajectories may thus have overlooked the rich repertoire of neuronal mechanisms (e.g., synaptic receptors or intrinsic conductances^18,19,29,30,38^). We therefore studied how synaptic and intrinsic factors interact to contribute to trajectory stability, controllability, and independence. We identified general trends: mechanisms that increase the separation between spontaneous and propagation regimes improve trajectory controllability, while mechanisms reducing downward transitions (from propagation to spontaneous activity) increase stability.

We further identified several biophysical ways to implement these reliability mechanisms. NMDA currents – known to maintain working memory (i.e., static attractor^61^) also stabilizes moving bump (dynamic attractor) propagation because of its slow, associative and positive feedback. Fast GABA-A currents limit unwanted initiation of spontaneous trajectories, and we predict that manipulating GABA-A time constant (by preserving the global inhibitory drive but perturbing inhibitory fluctuations) should specifically affect spontaneous replays. Frequency-independent inhibition-based mechanisms also enhance stability. This is consistent with mediodorsal thalamic activation, which increases fast spiking firing frequency in rodent mPFC, promoting stable neural sequences and working memory performance^6^.

Most notably, calcium-dependent CAN/AHP ionic conductances yielded the best combination of reliability (i.e., controllability, stability, and independence) and robustness. Our results overall suggest that spike-triggered ionic currents are essential for reliable attractors dynamics under noisy conditions. This aligns with previous suggestions that calcium-dependent conductances can support attractor dynamics, though these earlier relied on phenomenological descriptions^34,80^. Our findings generate experimental predictions, including whether calcium-dependent conductances mediate independent replay of multiple trajectories, as recently observed in the hippocampus^16^.

The large space of biophysical properties remains only partially explored, with potential roles for short-term plasticity^65^ or other intrinsic properties, such as rebound after inhibition, delayed dynamics due to slow potassium conductance, or intrinsic oscillations^29,30^. Our framework may also help interpret pathological dynamics. For example, schizophrenia involves unwanted spontaneous activity and unstable attractors^81^; attention-deficit hyperactivity disorder (ADHD) was proposed to arise from impaired gain modulation in central networks, which may compromise the stability of neural attractors^82^. Calcium-dependent conductances (CAN/AHP) might therefore constitute interesting targets in ADHD or schizophrenia.

Altogether, our results demonstrate the biophysical plausibility of reliable attractors under the presence of several mechanisms, in particular in the presence of calcium-dependent intrinsic currents. Their synergy with synaptic properties appears as a generic solution to regulate attractor reliability, depending on cognitive demands. Low levels of reliability are desirable because they prompt the self-generated emergence or clearance of representations required when exploration, creativity and cognitive flexibility are governing behavior. By contrast, higher levels of reliability are expected for exploitation, cognitive control, reliable decision-making and deterministic volition. The neuromodulation of the mechanisms examined here, in particular that of calcium-dependent intrinsic currents, might be essential to switch between these two general types of cognitive demands. Related, a major prediction of our study is that blocking the mechanisms considered (e.g., calcium-dependent intrinsic currents) in relevant neural structures (e.g., the PFC) should destabilise attractor dynamics observed experimentally^62,63^.

Finally, calcium-dependent intrinsic currents alleviate the reliance on synaptic connectivity alone and highlight the complementarity of synaptic (STDP) and intrinsic (CAN/AHP) properties. Such a combined solution would not have emerged from approaches that optimize the connectivity (artificial training) in networks of simplified “neuronal” units.

## Author contributions

MS JN BD designed the model and research project. MS DM JB BD simulated the model and analyzed its behavior. MS JN JB BD wrote the article.

## Funding

Centre national de la recherche scientifique (JN, BD)

Fondation pour la recherche médicale (JN)

Agence Nationale pour la Recherche (BD)

## Supplementary Figures

**Figure S1.**
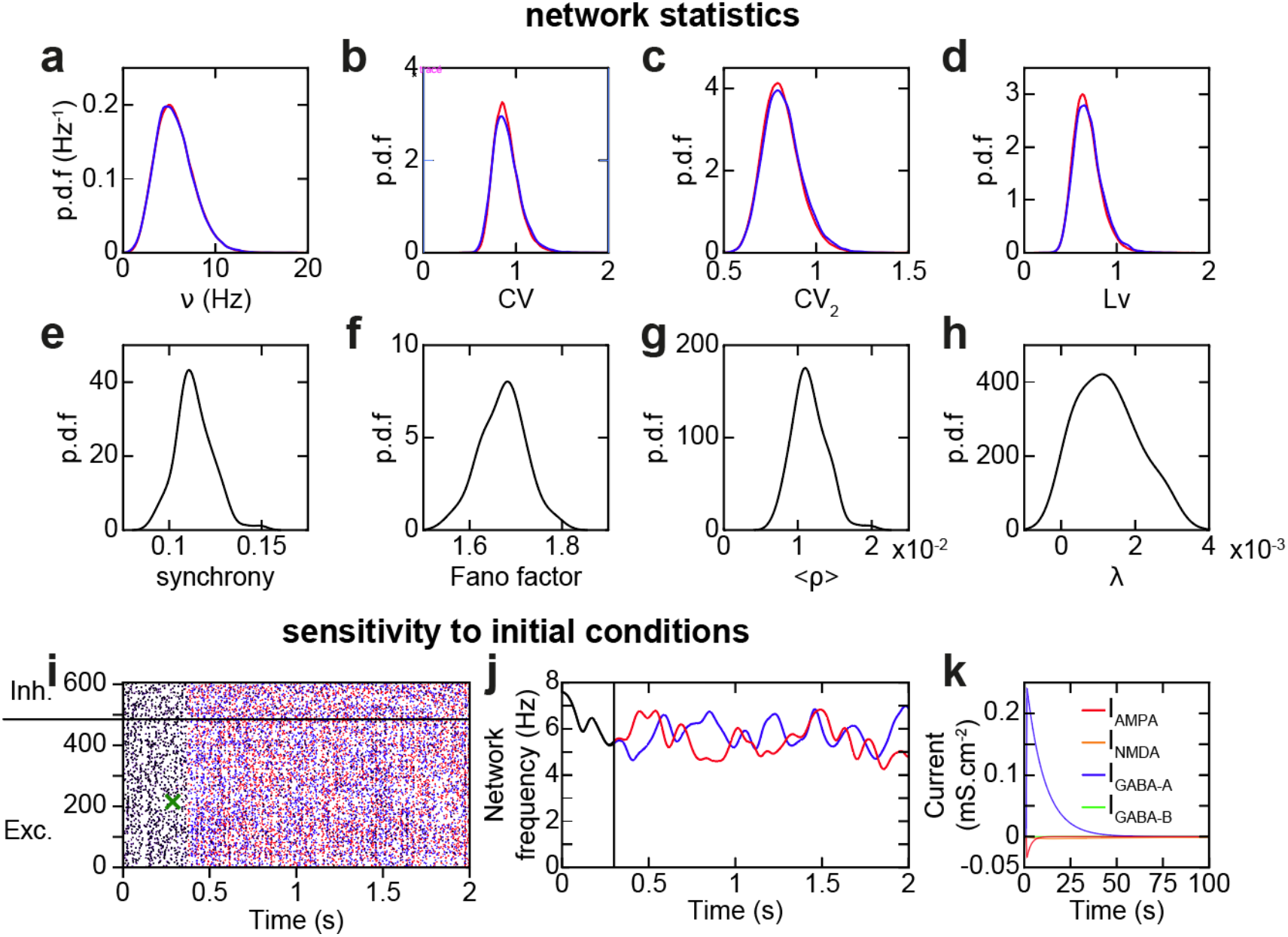
Distributions of network statistics over many network simulations. (**a-h**) Probability density functions of network spiking statistics, computed on 100 network simulations of 10s. Frequency (**a**), CV (**b**), CV2 (**c**), and Lv (**d**) of individual excitatory (red) and inhibitory (blue) neuronal spiking activity. Synchrony measure (**e**), Fano factor (**f**), average pairwise correlation coefficient (**g**), and Lyapunov exponent (**h**) of network spiking activity. (**i**) Chaotic network activity seen through sensitivity to initial conditions. A network was simulated in identical initial conditions, until a single randomly chosen spike at 300ms (green cross) was either kept (red spikes) or removed (blue spikes). Overlap in spikes between in both simulations are colored in black (notice that all spikes are identical and thus black before the green cross). (**j**) Same as (i), but average network frequency of both simulations (red & blue, overlap in black). (**k**) Stronger IPSC than EPSC balance total currents and thus fluctuation-based spiking. IPSC and EPSC are subdivided into their individual (AMPA, NMDA, GABA-A, GABA-B) components.

**Figure S2.**
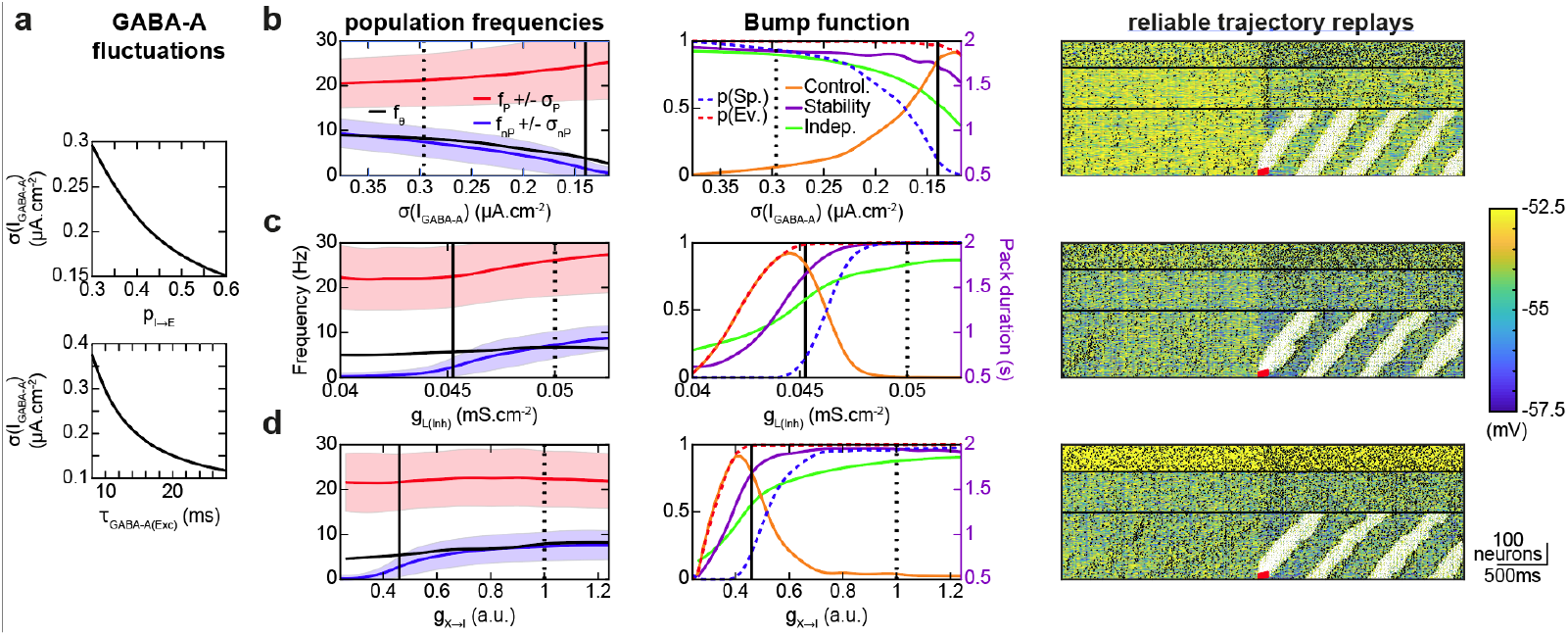
GABA-A fluctuations and alternative effects of lowering disinhibition on replay quality. (**a**) *σ*(*I*_*GABA-A*_) as a function of *f*_*I*→*E*_ or *τ*_*GABA-A*(*Exc*)_. (**b-d**) Same as **Fig. 5a-d**, but when varying the GABA-A current time constant of excitatory neurons *τ*_*GABA-A*(*Exc*)_ (*A*_*STDP*_ = 57.5) (**b**), leak conductance of inhibitory neurons *g*_*L,(Inh)*_ (*A*_*STDP*_ = 67.5) (**c**), and multiplicative factor *ρ*_*X*→*I*_ modulating the recurrent current conductances impinging upon inhibitory neurons *ρ*_*X*→*I*_ (*A*_*STDP*_ = 60) (**d**). For *τ*_*GABA-A*(*Exc*)_, *Δp* of inhibitory to excitatory neuron synapses were modulated in order for average *p*_*GABA-A*_ to be kept approximately constant at *v* = 5.5 *Hz*, i.e. weakened for longer *τ*_*GABA-A*(*Exc*)_. To do so, *Δp* was multiplied by the estimated average *p*_*GABA-A*_ value at *v* = 5.5 *Hz* for the standard value of *τ*_*GABA-A*(*Exc*)_ = 10 *ms* (computed as for 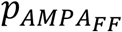) and divided by the same estimate but for the chosen value of *τ*_*GABA-A*(*Exc*)_.

**Figure S3.**
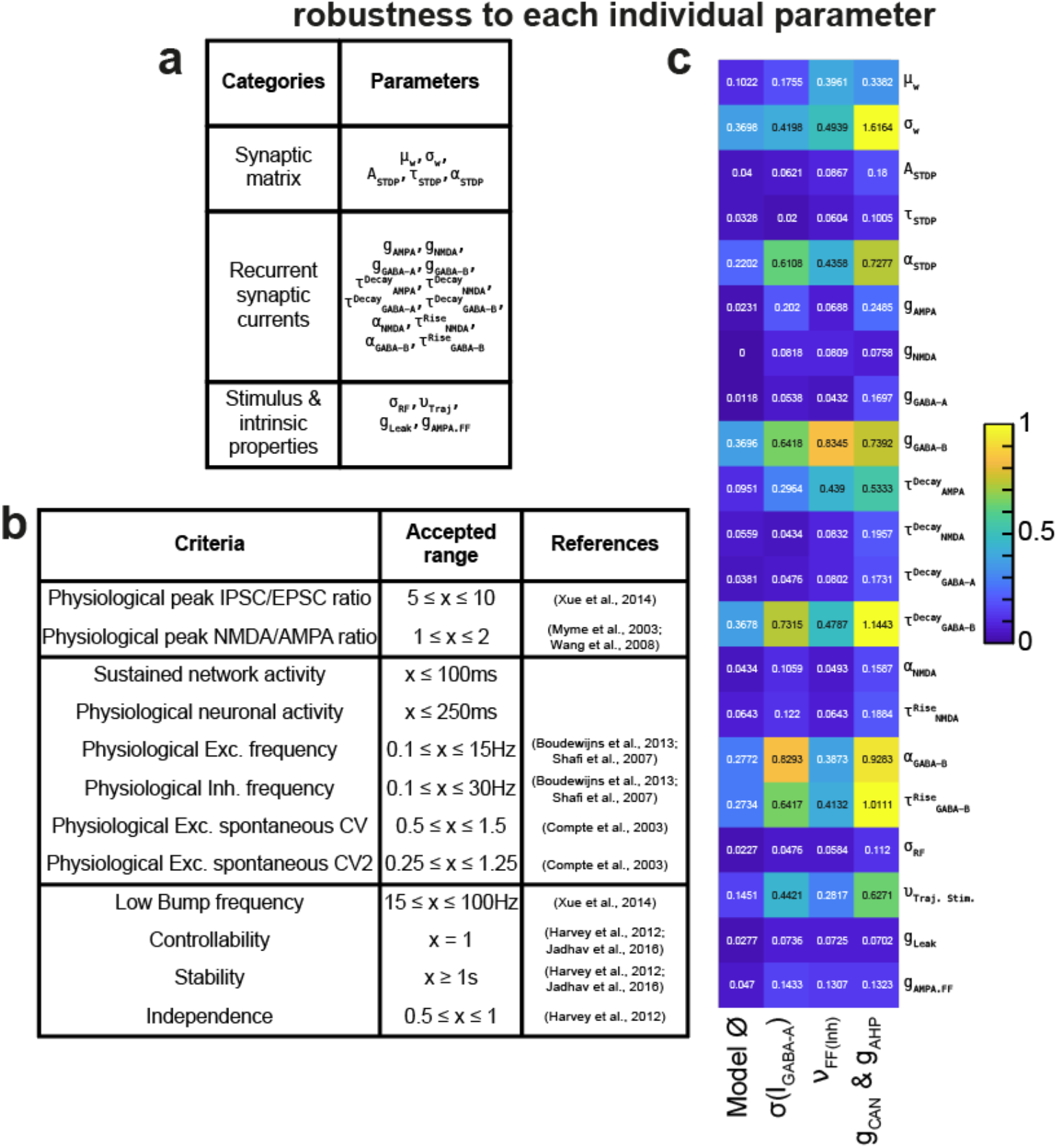
Robustness to parameters. Computation of the robustness score, quantifying to what extent the physiological low-frequency asynchronous irregular network activity with controllable, stable and independent trajectory replays is robust to the variation of the model’s parameters. (**a**) Parameters varied (see *Methods*). (**b**) List of criteria that need to be simultaneously met within a model network simulation for it to be considered biologically plausible. (**c**) Detail of the robustness score for each individual model parameter, for the different mechanisms and standard model (∅).

**Figure S4.**
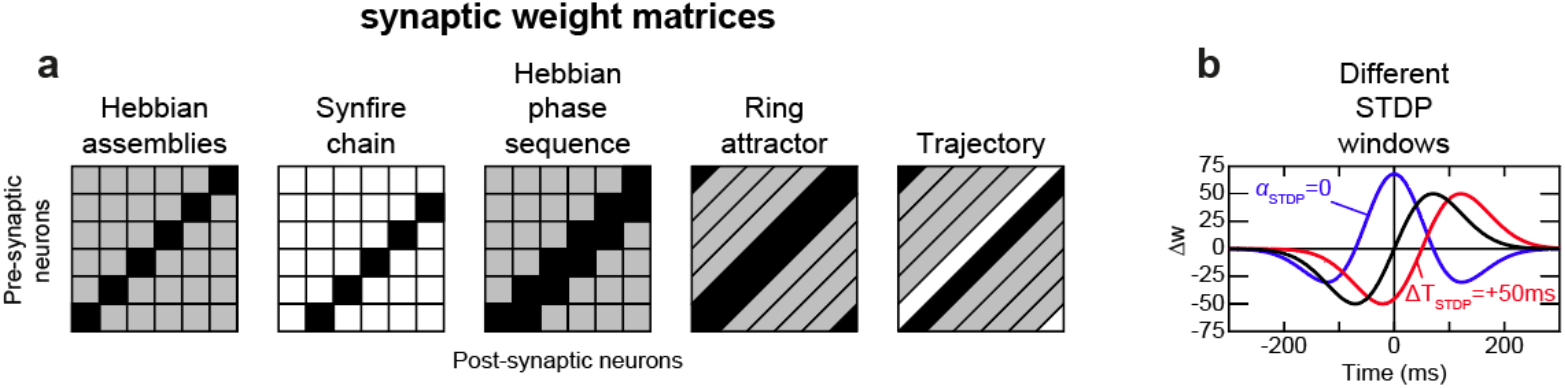
Synaptic weight matrices and STDP window parametrization for the control of static and dynamic discrete and continuous attractors shown in Fig. 7-9. (**a**) Weight matrices underlying the different types of network attractors. White, grey and black colors indicate the strength of synaptic weights (white = absence of synapses, grey = moderate weights, black = strong weights). (**b**) Modulation of the asymmetric STDP window (black) when varying its symmetry (*α*_*STDP*_ = 0, symmetric STDP window, blue curve) and temporal shift (*Δt* = +50ms, red curve).

## Methods

### Model of biophysical local recurrent neural network

We built a biophysical model of a generic local recurrent neural network, endowed with detailed biological properties of its neurons and connections, as in ^36^. The network model contained *N* neurons that were either excitatory (E) or inhibitory (I) (neurons projecting only glutamate or GABA, respectively ^84^), with probabilities *p*_*E*_ and *p*_*I*_ = 1 − *p*_*E*_ respectively, and 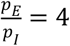. Connectivity was sparse (i.e. only a fraction of all possible connections exists, see *p*_*E*→*E*_, *p*_*E*→*I*_, *p*_*I*→*E*_, *p*_*I*→*I*_ parameter values ^41^) with no autapses (self-connections) and EE connections (from E to E neurons) drawn to ensure the over-representation of bidirectional connections in cortical networks (four times more than randomly drawn according to a Bernoulli scheme ^42^). The synaptic weights *w*_(*i,j*)_ of existing connections were drawn identically and independently from a log-normal distribution of parameters *μ*_*w*_ and *σ*_*w*_ ^42^. To cope with simulation times required for the massive explorations ran in the parameter space, neurons were modeled as leaky integrate-and-fire (LIF) neurons.

The membrane potential followed

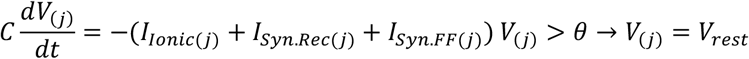

where neurons spike when the membrane potential reached the threshold *θ*, and repolarization to *V*_*rest*_ occurred after a refractory period *Δt*_*AP*_. Initial membrane potential of neurons were randomly drawn from a uniform distribution between *θ* and *V*_*rest*_.

The ionic current followed

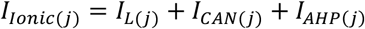

in which the leak current was

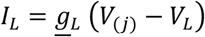

where *<u>g</u>*_*L*_ was the maximal conductance and *V*_*L*_ the equilibrium potential of the leak current.

The cationic non-selective (*I*_*CAN*_) current and the medium after-hyperpolarization (*I*_*AHP*_) currents, responsible for frequency adaptation and bistable discharge in pyramidal neurons, were taken as

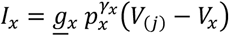

where *p*_*x*_ (*x* ∈ {CA*N, AHP*}) corresponded to the opening probability of both currents and *γ*_*x*_ the gating factor of opening probabilities. Denoting the intra-somatic calcium concentration as *Ca, p*_*x*_ followed

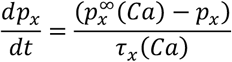

with

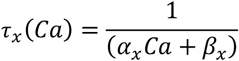

and

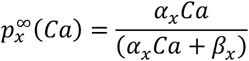

where *α*_*x*_ and *β*_*x*_ respectively denoted activation and deactivation kinetic constants, consistent with experimental data in layer 5 PFC pyramidal neurons ^43,44^.

The intra-somatic calcium concentration evolved according to discrete spike-induced increments and first-order exponential decay

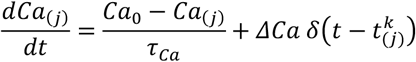

where 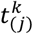 was the time of the *k*_th_ spike in the spike train of neuron *j, δ* the Dirac delta function, *τ* _*Ca*_ the time constant of calcium extrusion, *Ca*_0_ the basal calcium and *ΔCa* a spike-induced increment of calcium concentration.

The recurrent synaptic current on postsynaptic neuron *j*, from – either excitatory or inhibitory – presynaptic neurons (indexed by *i*), was

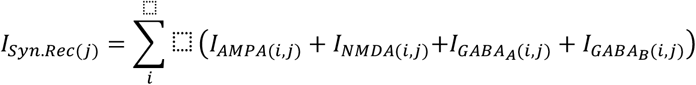

The delay for synaptic conduction and transmission, *Δt*_*syn*_, was considered uniform across the network ^39^. Synaptic recurrent currents followed

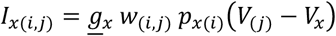

with *w*_(*i,j*)_ the synaptic weight. The NMDA current followed

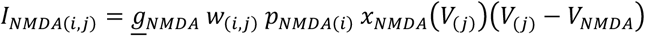

incorporating the magnesium block voltage-dependence modeled ^85^ as

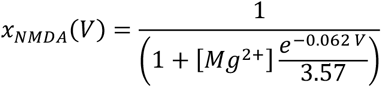

AMPA and GABA-A rise times were approximated as instantaneous ^39^ and bounded, with first-order decay

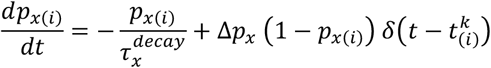

To take into account the longer NMDA and GABA-B ^86,87^ rise times, opening probabilities followed second-order dynamics ^39^

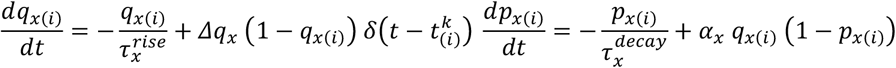

Recurrent excitatory and inhibitory currents were balanced in each postsynaptic neuron ^26^, according to driving forces and the excitation/inhibition weight ratio, through

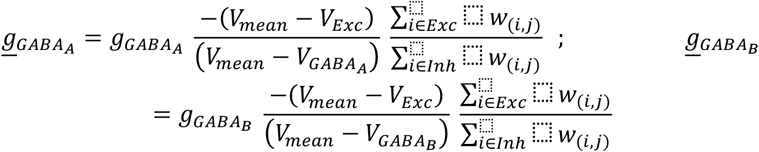

with 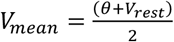 being an approximation of the average membrane potential. The excitation/inhibition weight ratio notably balanced the currents coming from inhibitory neurons with the 4x more numerous excitatory neurons (rendering inhibitory currents 4x stronger on average). When specified (**Fig. S2.d**), both excitatory and inhibitory conductances onto excitatory neurons were multiplied by *g* _*X*→*E*_, and onto inhibitory neurons by *g*_*X*→*I*_.

The feed-forward synaptic current *I*_*Syn*.*FF*(*j*)_ (putatively arising from subcortical and cortical inputs) consisted of an AMPA component

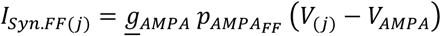

with a constant opening probability 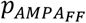, determined as the temporal average of AMPA channel openings due to *n*_*FF*_ neurons within putatively sub-cortical and cortico-cortical structures spiking at a given frequency *v*_*FF*_, following

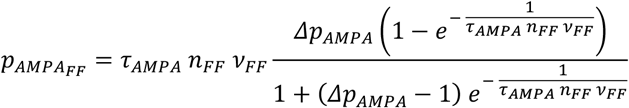

via integration (considering regular ISI for simplification during the integration). 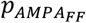 was considered constant so as to isolate the influence of deterministic chaos and spike irregularity on trajectory replay. However, to guarantee an initial stimulation sufficiently strong to start network activity, feedforward AMPA inputs were stronger at first (*n*_*FF*_ = 200 neurons, *v*_*FF*_ = 3 *Hz*) and progressively decreased during 250 *ms* to their final value (*v*_*FF*_ = 2.315 *Hz*; these initial 250 *ms* were cut from all figures and analyses). Trajectory replay was evoked 2s after the initial 250ms (**Fig. 2b-e**, red square) when the first 25 neurons of the trajectory received a strong feedforward AMPA stimulation (*n*_*FF*_ = 20 neurons, *v*_*FF*_ = *v*_*Traj*.*Stim*_ = 50 *Hz*, emulating a strong signal coming from a few neurons). The epoch before this trajectory-evoking stimulus was considered “Spontaneous” and the epoch after “Evoked”.

### Learning protocol

The neural network was subjected to “offline” learning, i.e. before the network simulation, during which the receptive fields of excitatory neurons were sequentially stimulated. The resulting neural frequency conditioned learning of synaptic weights via STDP between excitatory neurons. This “offline” learning procedure would correspond to the trajectory stimulus being learned and memorized long before the network simulation.

Neuronal receptor fields existed in a 2D spatial area (**Fig. 2a** left) following non-normalized bivariate Gaussian functions around their center points (*x*_*j*_, *y*_*j*_) organized along a square grid. For a stimulation point *s* (*x*_*s*_(*t*_*s*_), *y*_*s*_(*t*_*s*_)) of intensity *I*_*s*_ at moment *t*_*s*_, the resulting neural frequency of the stimulation of the receptive field was

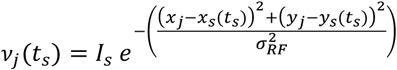

This stimulation was part of a dynamic spatiotemporal trajectory moving as time went by. The synaptic weights between neurons were then altered in proportion to their frequencies according to a phenomenological STDP rule (see below). A circular trajectory was chosen in order to study the sequence replay stability across multiple circle loops. The trajectory stimulus advanced by 0.05 (in the spatial area reference) every *dt*_*Traj*_ = 20 *ms* time step, with a small overlap between the trajectory start and end to ensure looping. For discrete stimuli (**Fig. 5**), the trajectory cycled 10x through the shown sequence of black dots (with the same trajectory time step). Neurons were considered as belonging to the trajectory when any of their stimulation-induced *v*_*j*_ (*t*_*s*_) > 5% of the maximum neuron frequency the trajectory produced.

### Spike-timing dependent plasticity

We assessed various STDP temporal windows, from entirely asymmetric (*α* _*STDP*_ = 1) to symmetric (*α* _*STDP*_ = 0) and time-shifted (*ΔT*_*STDP*_) functions. To modulate STDP symmetry, we identified two STDP functions, an asymmetric and a symmetric one (whose integrals equal 0, so that LTP and LTD contributions are balanced), and then performed a linear combination of both to obtain various degrees of STDP temporal asymmetry. However, even though the integral stayed null, the integral of the positive part changed, which we corrected by normalizing according to the asymmetric function’s integral’s positive part. As such, the STDP function followed

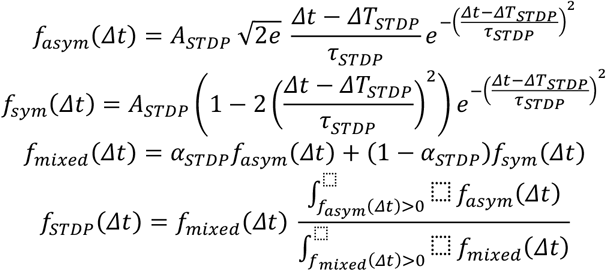

where *Δt* = *t*_*post*_ − *t*_*pre*_ was the temporal difference between pre- and postsynaptic spikes, *A*_*STDP*_ the STDP amplitude and *τ* _*STDP*_ the STDP time constant. As such, taking into account the frequencies of pre- and postsynaptic neurons and the time difference between stimulation times, the weights were changed according to

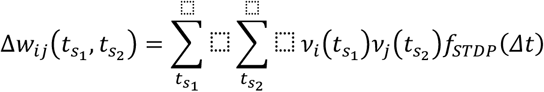

with 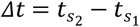. A lower hardbound limit (*w* ≥ 0) was imposed after STDP learning, whereas no upper hardbound limit was imposed. The respective firing frequencies of the populations are taken into account as it has been shown that STDP is essentially dependent upon firing rate rather than spike timing under natural conditions, i.e irregular spiking ^88^. The description employed here directly reflects the multiplicative dependance of synaptic modifications upon presynaptic and postsynaptic firing rates, modeled in a more detailed fashion (with calcium-dependent kinases and phosphatases) in our previous study ^36^.

### Synaptic scaling

In order to keep neuronal activity within certain putative homeostatic bounds, synaptic weights entering a postsynaptic neuron are subjected after STDP learning to a simple multiplicative phenomenological form of synaptic scaling ^51^, potentially representing hetero-synaptic LTD, where the sum of weights impinging upon a pyramidal neuron is kept constant before and after STDP. This is written

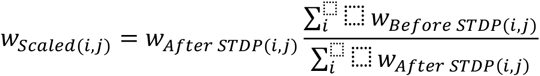

### Detection of bumps

In order to detect propagating activity bump along the synaptic pathway, we first convolved neural spiking activity with a centered normalized Gaussian function where *σ* = 30*ms*, to then spatially convolute it with the bivariate Gaussian receptive field function (see above) centered on the discrete points of the spatiotemporal trajectory. Such smoothing procedures allowed us to reliably choose a frequency threshold (12.5*Hz*) above which trajectory points were considered “active”. Conversely, from these “active” trajectory points, we considered trajectory neurons “active” when at least 40% of the trajectory points having stimulated that neuron’s receptive field (above the aforementioned 5% of maximum neuron frequency), weighted by the neural frequency resulting from trajectory stimulation, were “active”. This allowed us to define bump emergence as when at least 20 dynamically changing trajectory neurons were “active” on average during 500 successive milliseconds (ensuring activity packets were strong enough, e.g. **Fig. 2b-e** white spikes).

### Determining bump and non-bump frequency average, fluctuations or threshold

*f*_*nB*_ and *σ*_*nB*_ were determined as the frequency average and fluctuations of the aforementioned spatially-convoluted trajectory points of neurons outside the bump during periods without bumps, while *f*_*B*_ and *σ* _*B*_ were similarly determined but for neurons within the bump during bumps. These frequency averages were done across neurons and network realizations. By manipulating the frequency of neurons within the trajectory *f*_*T*_ through different levels of feedforward AMPA currents, the frequency threshold *f*_*θ*_ ms determined as the minimal *f*_*nB*_ for which bumps propagate constantly (≥ 1900 *ms* out of 2 *s* total, and *p*_*Spont*_ ≥ 0.95). This understanding derived from the predictions of the bistable and noisy regime transition model, mimicking the process where the non-bump frequency *f*_*nB*_ stable fixed point increases until it coalesces with the threshold *f*_*θ*_ unstable fixed point (as in a saddle-node bifurcation), in which case only the bump frequency *f*_*B*_ stable fixed point remains and bumps thus propagate constantly. As a side remark, studying the role of intrinsic biophysical mechanisms being the aim of the present study, using frequency observables as described above does not imply that our findings could have been reproduced with a rate model.

### Maximum Lyapunov Estimate

To quantify the chaotic nature of the network’s activity, we estimated the maximum Lyapunov exponent *λ* on the one-dimensional time series of the estimated instantaneous spiking frequency (*σ* = 30 *ms*) averaged across excitatory neurons ^89^. To do so, we reconstructed the phase space through time-delay embedding with heuristics agreed upon in the literature ^90,91^. The lag length was estimated as the first lag length for which the autocorrelation coefficient *AC* < *e*^1 92^. The embedding dimension was estimated via a MATLAB program developed by Mirwais Kizilkaya according to the false nearest neighbor method ^92,93^ as the minimal dimension with 0% false nearest neighbors as determined by tolerance factors (*R*_*tol*_ = 10, *A*_*tol*_ = 2, ^66^).

### Spiking variability and synchrony

Spiking variability and synchrony measures are computed as in ^36^. To compare spike variability between our model and experimental data, we quantified the coefficient of variation (CV) of the inter-spike interval (ISI) distribution of the spiking trains of neurons in the network ^46^ according to

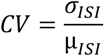

However, the CV measure assumes stationarity of the data. Since this assumption is not necessarily verified, we also computed the CV2 and Lv of the spike trains to evaluate the variability of ISIs at a local level, according to

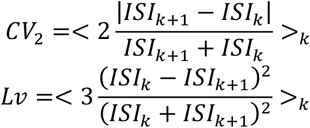

CV = *CV*_2_ = *Lv* = 1 for an ISI distribution drawn from homogeneous Poisson spike trains and = 0 for perfectly regular spike trains (all ISI are equal). *CV* typically stand around 1 to 1.5 *in vivo*, while CV2 and Lv stand around 0.25 to 1.25 and 0 to 2 respectively *in vivo* ^46^. CV was computed on all ISI, while CV2 and Lv are computed for each neuron then averaged across neurons.

Multiple synchrony measures were computed ^47,94^, a synchrony measure *S*, pairwise correlation coefficient averaged over all pairs of neurons < *ρ* >, and Fano factor *F*, following

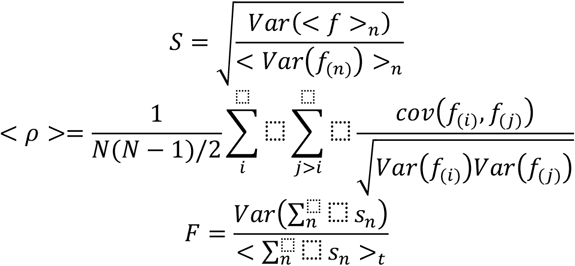

where *f* was the estimated instantaneous neural spiking frequency via Gaussian convolution (*σ* = 30*ms*), *n* the neuron index, and *s* the population sum of spike counts, where 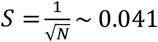, < *ρ* >= 0 and *F* = 1 for perfectly asynchronous network activity, and *S* =< *ρ* >= 1 while F increases for perfectly synchronous network activity.

### Protocol for assessing the nature of intrinsic bistability

The protocol for evaluating the nature of neural intrinsic bistability, taken from ^38^, consisted of a strong phasic input (of amplitude 2*θ*_*on*_ during 200 *ms*) followed by a weaker delay-period tonic input (of amplitude *I*_*Inj*_ during 10*s*), in order to reveal conditional bistability activated by the phasic input but conditional on (i.e. requiring the) weaker delay-period tonic input. *θ* _*on*_ corresponded to the minimal delay-period tonic input current required to induce sustained firing during the delay without the strong initial phasic input, and *θ* _*off*_ the same but with the strong initial phasic input. *θ* _*transient*_ was the same as *θ* _*off*_ but corresponded to the minimal delay-period tonic input required to induce unstable (rather than sustained) firing. Firing was considered sustained when there were three or more spikes during the last 2 *s* of the tonic input with stable ISIs (determined when 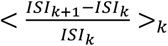 was inferior to 0.05). Otherwise, firing was considered unstable for a single spike beyond 25ms after the initial phasic input, or for two spikes or more during non-sustained firing.

When *θ* _*on*_ = *θ*_*off*_ = *θ*_*transient*_, the neuron was considered monostable, i.e. the strong initial input current did not activate any intrinsic mechanisms generating sustained firing. When *θ* _*on*_ = *θ*_*off*_ > *θ*_*transient*_, the neuron was considered transiently bistable, the strong initial input inducing weak mechanisms generating unstable (but not sustained) firing. When *θ*_*on*_ > *θ*_*off*_, the neuron was considered conditionally bistable, since the delay-period input, weaker than the initial phasic input but non-zero, could induce sustained firing, bistability being thus conditional upon the delay-period input. Finally, if *θ*_*on*_ > 0 > *θ*_*off*_, the neuron’s bistability was considered absolute, i.e. sustained neuronal firing after an initial input lasts until a hyperpolarizing current stops it.

### Estimating robustness to variability of the model’s parameters

We studied how sensitive the phenomenon of interest (namely controlled, stable and independent trajectory replay with asynchronous irregular network dynamics) was to the variability of model parameters, since biological systems present strong variability. To do so, we systematically varied important parameters, and defined a list of criteria which all need to be met (**Fig. S3**), encompassing physiological peak conductance ratios (top row), spiking activity regime (middle row) and controllable stable independent trajectory replays (bottom row). Sustained network activity (middle row) was determined when the maximal duration without network spikes was 100ms, to exclude strongly oscillating networks prohibiting controllable trajectory replay. Physiological neuronal activity was determined when neuronal activity was 100Hz at most for 250ms (in order to exclude trivial trajectory replay cases where replay was actually detected as a single neuron stably firing at 100Hz during 500ms). CV and CV2 were determined during the spontaneous epoch (before the trajectory replay evoking stimulus at 2s, **Fig. 2b-e**).

Parameters were varied over a range of 40 equally-spaced values, generally spanning 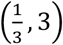 x the standard parameter value. Network simulations were repeated 5 times for each value (due to the potential variability of trajectory replays), with each repetition being evaluated independently concerning the criteria. The robustness score was computed as

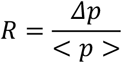

where < *p* > represented the average correct parameter value (weighted by the proportion of correct repetitions), and *Δp* the sum of correct parameter steps (once again weighted by the proportion of correct repetitions), where a step was the difference between the next and previous parameter value divided by 2, or 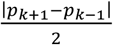 (values being equally spaced).

This robustness score was conservative no matter the arbitrarily chosen range, since it is a biased underestimation which approaches its true maximal limit value given an optimal chosen range. Indeed, robustness was limited by 1) how large the parameter range considered was, and 2) how close parameter steps were. The robustness score decreased from its true maximal value with ranges which were too small and step values too large. Contributions of individual criteria (**Fig. 6c**, right) were computed as the *Δp* when considering only that one criteria (with the same < *p* > value still computed over all criteria, for better comparison of individual contributions).

### Numerical integration and parameters of the biophysical network model

Models were simulated and explored using custom developed code under MATLAB and were numerically integrated using the forward Euler method with time-step *Δt* = 0.5*ms* in network models. The code MATLAB (tested on 2018b) is provided along the article. Reproducing precisely some of the figures requires loading the corresponding biophysical parameters that were deposited here: https://datadryad.org/stash/share/i_Mz55853ZGXvi6Q1qUdLWDJtHeDqbb4sfLRTDkZP2U.

Unless indicated in figure legends, standard parameter values were as following. Concerning the network architecture, *N* = *n*_*Exc*_ + *n*_*Inh*_ = 605 neurons, *p*_*Exc*_ = 0.8, so that *n*_*Exc*_ = *N, p*_*Exc*_ = 484 neurons and *n*_*Inh*_ = *N, p*_*Inh*_ = 121 neurons. Concerning Integrate-and-Fire neuron properties, *C* = 1 *μF. cm*^−2^, *V*_*rest*_ = −65 *mV, θ* = −50 *mV*, 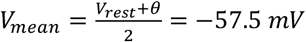, *Δt*_*AP*_ = 3 *ms*. Concerning ionic currents, *<u>g</u>*_*L*_ = 0.05 *mS. cm*^−2^, *V*_*L*_ = −70 *m*V, <u>g</u>_*CAN*_ = 0 *mS. cm*^−2^, *V*_*CAN*_ = 30 *mV, α*_*CAN*_ = 0.03125 *μM*^−1^. *ms*^−1^, *β*_*CAN*_ = 0.025 *ms*^−1^, *γ*_*CAN*_ = 1, *<u>g</u>*_*AHP*_ = 0 *mS. cm*^−2^, *V*_*AHP*_ = −90 *m*V, *α*_*AHP*_ = 0.125 *μM*^−1^. *ms*^−1^, *β*_*AHP*_ = 0.025 *ms*^−1^, *γ*_*AHP*_ = 1, *ΔCa* = 0.2 *μM, Ca*_0_ = 0.1 *μM, τ*_*Ca*_ = 100 *ms*. Concerning the weight matrix, *μ*_*w*_= 0.03, *σ*_*w*_ = 0.015, *p*_*E*→*E*_ = *p*_*E*→*I*_ = *p*_*I*→*I*_ = *p*_*I*→*E*_ = 0.3. Concerning synaptic currents, 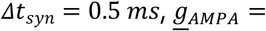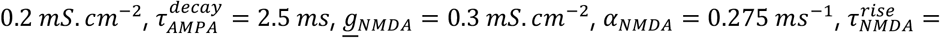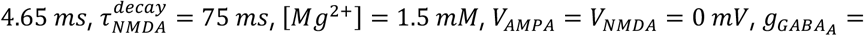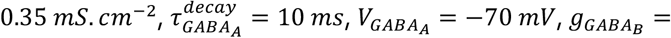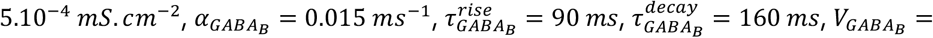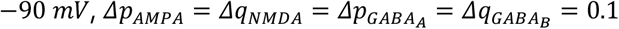 . Concerning the learning procedure and STDP, *σ*_*RF*_ = 0.13, *I*_*S*_ = 0.02925, *τ*_*STDP*_ = 100 *ms, A*_*STDP*_ = 50, *α*_*STDP*_ = 1, *ΔT*_*STDP*_ = 0 *ms*.

Parameters for the biophysical mechanisms (**Fig. 4**) were systematically determined as the (*A*_*STDP*_, mechanism parameter) value couple maximizing the product of controllability, stability and independence (all three being normalized between 0 and 1).

For the bidirectional ring attractor (**Fig. 5.e2**), model parameters were as followed: *p*_*I*→*E*_ = 0.4, *g*_*X*→*E*_ = 1.5, *g*_*X*→*I*_ = 0.5, *<u>g</u>*_*AHP*_ = 0.2 *mS. cm*^−2^, *α*_*AHP*_ = 0.001 *μM*^−1^. *ms*^−1^, *β*_*AHP*_ = 0.002 *ms*^−1^, *γ*_*AHP*_ = 2, *I*_*S*_ = 0.03.

### Reduced model of bump dynamics

We developed a reduced version of the full model to catch essential features of the regime of a bump of activity between neurons within the engram. Whereas noisy spiking sets asynchronous-irregular global dynamics, we found that the simplest observable distinguishing bump from non-bump neurons was their firing frequency. We therefore based our theoretical analysis – (see the following two sections, Propagation condition model and Regime transition model) on a rate-based simplified description of collective neuronal dynamics, which also incorporated the effect of noisy fluctuations on bumps. This analysis allowed us to derive qualitative predictions regarding the effects of biophysical parameters on bump propagation and transitions, that were in qualitative accord with full model simulations (see below and *Results*). Frequency parameters of the reduced model were estimated from neural activity in simulations of the simpler version, with no additional biophysical mechanism of the full recurrent network model (the standard “Model ∅”). The reduced model was thus neither devised as a specific quantitative tool (see the text corresponding to Fig.4c), nor compared quantitatively against the full parameter space of the whole model.

### Propagation condition model

Basically, propagation requires that, on average, spiking at frequency *f* in (upstream) presynaptic neurons must induce spiking at a frequency superior or equal to *f* in (down-stream) postsynaptic neurons. Therefore, we wrote a set of equations where presynaptic AMPA and NMDA input currents to a postsynaptic neuron are scaled by the firing frequency *f* of the presynaptic neuron and searched for frequency conditions where postsynaptic neurons fire at a frequency greater or equal than *f*. This *propagation condition model* is an extremely simplified one-dimensional reduced representation of bump propagation within the local cortical recurrent network. This model is space-free and shall be considered as a representation of internal dynamics within the bump during its propagation, i.e. in a referential moving at the speed of bump propagation. Noticeably, the propagation condition model only considers a pre-/postsynaptic feedforward interaction, but does not take into account possible recurrent effects of the postsynaptic neuron on the presynaptic neurons or on the network. The propagation condition model nevertheless considers incoming excitatory and fixed inhibitory inputs from the entire network onto the postsynaptic neuron. These inputs are lumped together into common AMPA, NMDA and GABA-A terms that are quantitatively fitted on average synaptic currents impinging bump neurons in network simulations. The additional assumption is made that excitatory currents are essentially provided by upstream neurons within the bump (vs from neurons outside the bump, whether inside or outside the trajectory), so that AMPA and NMDA currents are scaled by *f*.

To track the problem in a deterministic way, we leveraged from the observation that, regardless of whether neurons spike in the spontaneous regime or within the bump, 1) ISIs generally terminate through rapid final depolarizing fluctuations, due to chaotic dynamics, and that 2) these fluctuations start *Δt*_*fluct*_∼15*ms* before spiking. We numerically determined, from all ISIs during bump activity in network simulations, the mean time-to-spiking *Δt*_*spiking*_(*V, f*), as a function of the membrane potential and the firing frequency of the current ISI. We found that *Δt*_*spiking*_(*V, f*) = *Δt*_*fluct*_ was largely independent of firing frequency, which allowed us to numerically estimate 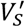 (around -53 mV).

We also considered, based on neuronal dynamics in the network model, that the membrane potential was essentially deterministically driven – before reaching 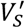 and the final fluctuation to spiking – by average input and leak currents. Thus the membrane potential converged exponentially to its steady state *V*^∗^ with

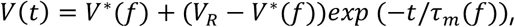

where *V*^∗^(f) was obtained from the equilibrium of ionic currents at steady-state

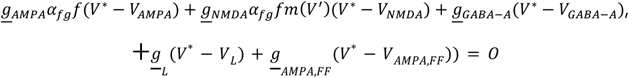

with *α*_*fg*_ a conversion factor for dimensional compatibility and the non-linearity of the NMDA approximated to its value *V*^′^, such that one can solve explicitly in terms of *V*^∗^:

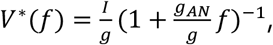

with

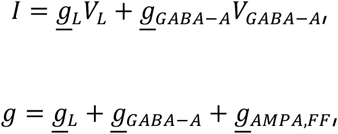

and

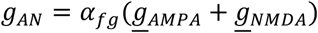

which could be linearized 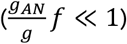 to

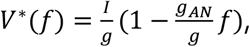

the membrane time-constant being written

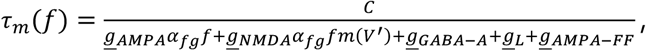

and

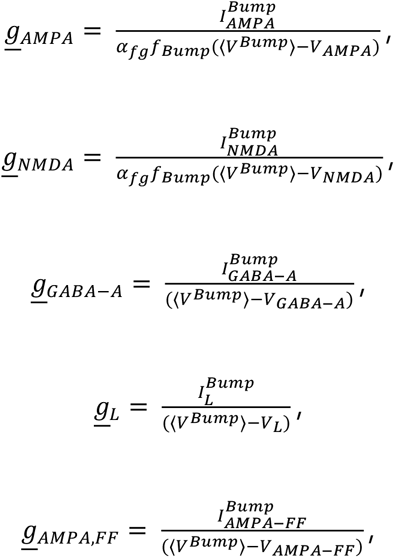

being estimated from bump mean membrane potential ⟨*V*^*Bump*^⟩, mean currents (see below) and mean firing frequency *f*_*Bump*_ obtained from network simulations.

As a final step of the propagation condition model, we then computed:

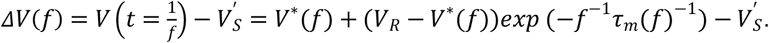

A negative *ΔV*(*f*) indicates that the potential has not yet reached, at time 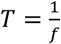, the threshold 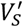 yielding rapid fluctuation-driven spiking so that postsynaptic frequency is lower than *f*, the presynaptic firing frequency. Therefore, propagation fails. Conversely, a positive *ΔV*(*f*) indicates that the postsynaptic frequency exceeded the presynaptic one, so that propagation continues downstream. Finding a critical frequency *f*_*θ*_ such that *ΔV*(*f*_*θ*_) = 0 indicates that pre- and postsynaptic frequencies are equal and propagation of spiking occurs at frequency *f*_*θ*_. Moreover, the slope of *ΔV*(*f*) at *f*_*θ*_ determines the stability of the propagation. A negative slope indicates a stable propagation at frequency *f*_*θ*_ as fluctuations (due to chaotic network dynamics) will be quenched out by restoring forces driving the frequency back to *f*_*θ*_ (frequency increases below *f*_*θ*_ (*ΔV*(*f*) > 0) and decreases above it (*ΔV*(*f*) < 0)). A positive slope, to the contrary, indicates an unstable propagation with firing frequency ineluctably diverging from *f*_*θ*_.

Computing the model indicated that, under our simplifying hypotheses, a single critical frequency *f*_*θ*_ was found at which the slope of the *ΔV*(*f*) was positive (see Results). Therefore, the propagation condition model suggested that *f*_*θ*_ corresponded to an unstable fixed-point in the frequency dimension, acting as a threshold that separated, for trajectory neurons, the spontaneous regime (no bump propagation) from the regime of bump propagation. Actually, the propagation condition model predicted the value of *f*_*θ*_ quite well (see *Results*), with a value very close to that directly estimated from network simulations (see below). The quality of the propagation condition model was also evaluated by computing mean currents and comparing them to those found in network simulations (see *Results*). Currents were computed as:

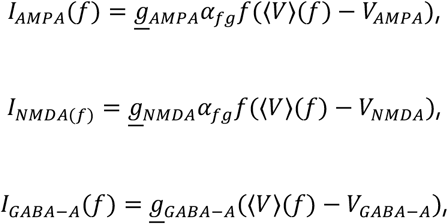

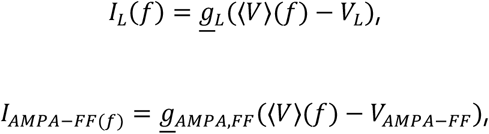

Where

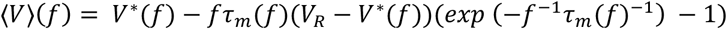

### Regime transition model

Although *f*_*θ*_ and the average currents (see *Results*) underlying the propagation condition were estimated, the model was however unable to identify the two stable frequency fixed-points *f*_*non-Bump*_ and *f*_*Bump*_ setting the average spiking frequency in the spontaneous regime and during bump propagation in network simulations. This was because the simplifications regarding recurrent interactions within the network between excitatory neurons within the bump and neurons outside the bump (i.e. excitatory neurons inside and outside the trajectory, and inhibitory neurons) were too strong to account for the non-linearity ensuring negative feedbacks in the vicinity of *f*_*non-Bump*_ and *f*_*Bump*_ stable fixed-points.

However, to better understand propagation of the bump within the network, we considered the co-existence of the unstable *f*_*θ*_ fixed-point and of the two stable *f*_*non-Bump*_ and *f*_*Bump*_ stable fixed-points to build a phenomenological one-dimensional reduced *regime transition* model. Moreover, to evaluate the ability of this simplified model in explaining complex propagation behavior in the whole network by a simple model based on an unstable fixed-point separating two spontaneous and bump propagation regimes, we included a stochastic component and determined to which extent the simplified propagation model was able to account for transition rates between the spontaneous and propagation regimes in trajectory neurons. Specifically, the probability of the emergence of propagating bumps from the spontaneous regime, *p* (*Spont*. ), the probability of propagating evoked bumps *p*(*Evoked*) and their duration was computed from the model.

In the model, the firing frequency of neurons within the trajectory followed:

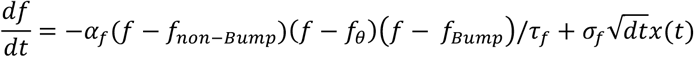

where *α*_*f*_ is a scaling factor, *τ*_*f*_ = *τ*_*m*_(*f*_*θ*_) (see above), *x*(*t*) is a Gaussian stochastic variable with mean 0 and standard deviation 1 and *σ*_*f*_= *σ*_*non-Bump*_ for *f* < *f*_*criterion*_ and *σ*_*f*_ = *σ*_*Bump*_ for *f* ≥ *f*_criter*i*on_ with *σ*_*non-Bump*_ and *σ*_*Bump*_ estimated from network simulations. The empirical estimation of *f*_*θ*_ in the network model was obtained by finding the frequency best separating Bump and non-Bump frequency distributions (see above). The noise, accounting for stochastic state transitions, is white. This choice is very classical and not essential to our results . The *α*_*f*_ parameter allowed us to fit the order of magnitude of transition rates. Its exact value has no impact on the qualitative interpretation of mechanisms. Frequency parameters of the theoretical models were estimated only for the standard model (“model ∅”).

### Parameters

*α*_*fg*_ = 1*mS. cm*^−2^. *Hz*^−1^, 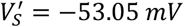, ⟨*V*^*Bump*^⟩ = −53.23*m*V, *f*_*non-Bump*;_ = 6.48 *Hz, f*_*Bump*_ = 14.34 *Hz, f*_*θ*_ = 9.7 *Hz, α*_*f*_ = 2*b* − 3 *s*^2^, *σ*_*non-Bump*_ = 1.96 *Hz, σ*_*Bump*_ = 3.55 *Hz*.

